# Mechanoscribed Xolography for Multiscale Mechanical Patterning of Cell-laden Hydrogels

**DOI:** 10.64898/2026.10.18.759929

**Authors:** Lena Elliger, Tom C. van Hattem, Mariana A. G. Oliva, Tess. L. van Alphen, Niklas F. König, Paul Genever, Sandra Hofmann, Keita Ito, Lorenzo Albertazzi, Manuel Salmeron-Sanchez, Miguel Castilho

**Author notes:** These authors contributed equally to this work (Lena Elliger, Tom C. van Hattem).

## Abstract

Mechanotransduction is a key regulator of healthy and malignant tissue development. However, current mechanobiological insights are largely derived from cells cultured on two-dimensional substrates or in homogeneous biomaterials, which insufficiently capture the three-dimensional (3D) heterogeneity and dynamic nature of the native extracellular matrix. This gap persists due to a lack of manufacturing processes and analysis modalities to control and monitor material properties and their evolution in 3D.

Here, Mechanoscribed Xolography is introduced as a light sheet volumetric bioprinting process to rapidly fabricate 3D cell-laden hydrogels with spatially controlled mechanical heterogeneity across multiple length scales. Moreover, synchronizing in-situ optical microscopy with printing enables autonomous detection and spatial location-driven printing around cellular spheroids.

The refinement and spatiotemporal evolution of material properties are assessed with Brillouin microscopy, which verifies that the contrast of imprinted heterogeneity is preserved over three weeks. Remarkably, regional differences in cell behavior are attained within a one-step, one-material construct with elasticity of approximately 0.3 kPa–0.8 kPa, or 0.7 kPa–2.9 kPa with a crosslinking-enhancing additive. This foundational work on high-precision, rapid bioprinting surrounding cellular objects represents a unique tool to systematically advance experimentation of 3D mechanotransduction and prospectively engineer cell-instructive biomaterials for regenerative medicine.

## 1. Introduction

Biological tissues are heterogeneous structures whose mechanical properties evolve over time and space.^1^ Cells embedded in the extracellular matrix (ECM) sense these properties through mechanotransduction, and both elasticity and viscoelasticity of the ECM have been shown to influence morphology, fate commitment, and matrix remodeling.^2–5^ Not only the absolute values of these mechanical cues, but also their spatial organization regulates cellular response: the distribution of mechanical heterogeneity is, for instance, shown to influence the expression of osteogenic differentiation markers.^4,6–8^ Importantly, because this mechanical heterogeneity is perceived by individual cells, it is relevant from the nanoscale to microscale, and may differ from the macroscopically observed heterogeneity on the tissue level. Moreover, most of what is known about mechanotransduction derives from two-dimensional substrates or homogeneous biomaterials and has limited transferability to the heterogeneous 3D tissue environment.^2,9–12^ This gap originates from two constraints: engineering mechanical heterogeneity within soft, cell-laden 3D hydrogels at the scale of cells, and monitoring its evolution non-destructively over the culture period.

Light-based volumetric additive manufacturing (VAM) addresses the first constraint by modulating energy input within a volume, generating spatially distinct degrees of crosslinking. Most 3D approaches employ this concept for the controlled placement or immobilization of bioactive molecules.^13,14^ Few technologies, like Filament Light (FLight) biofabrication, control spatial mechanics. FLight creates aligned microfilaments within a photoresponsive hydrogel by utilizing light beam speckle patterns and self-focusing effects during crosslinking, which produce mechanical anisotropy that aligns chondrocytes and, more recently, myoblasts.^12,15,16^ Because the filament growth propagates unidirectionally throughout the photoresin, voxel-specific control of mechanical properties and with this targeted control of the environment of individual cells, is constrained. Xolography instead confines polymerization to the intersection of two wavelengths by requiring quasi-simultaneous exposure of a dual-color photoinitiator (DCPI) to both.^17^ Unlike tomographic approaches such as computed axial lithography (CAL), it does not rely on cumulative energy doses to reach a photopolymerization threshold, which minimizes background curing.^18,19^ Moreover, the second light dose can be modulated locally and independently, introduced as grayscaled Xolography, whereas in CAL any local adjustment of the energy dose inevitably affects other regions within the projection paths.^20^

Grayscaled Xolography has been demonstrated with low molecular-weight polyethylene glycol diacrylate (PEGDA), though this formulation lacks cytocompatibility.^20^ Gelatin methacryloyl (GelMA)-based formulations have since been grayscaled, but cytotoxicity associated with the coinitiator triethanolamine (TEOA) and the lack of cellular readouts leave the biological relevance of mechanical patterns questionable.^21,22^ Cytocompatible GelMA formulations have been separately improved through an updated DCPI, BIS-TRIS as coinitiator, and refractive index matching during printing.^21–23^ However, grayscaled bioprinting of cell-laden photoresins has nonetheless not been achieved, and it remains unknown whether distinct crosslinking states close to cellular scale through grayscaling can produce corresponding regional cellular response.

An additional challenge is specific to cell-laden VAM: Cells and spheroids are randomly distributed within the photoresin and cannot be positioned, so a defined environment can only be assigned by adapting the printed geometry to their actual location, which requires their detection in the resin. In-situ monitoring has previously assessed print accuracy through Schlieren refractive index mapping and transmission imaging, in which corrections were fed into subsequent prints, as well as real-time imaging during printing.^24–26^ However, monitoring only informed the dose intensity, never the printed geometry itself. Position-registered printing in CAL adapts print geometry to the position of objects, but is dependent on fluorescent labeling of the objects and constrained by the spatial coupling of cumulative energy doses described above.^27^

The second constraint concerns 3D characterization of heterogeneous properties. Nanoindentation or atomic force microscopy are able to resolve features on or below the cellular level, however, they are surface-bound techniques that cannot probe 3D mechanical properties. Recently, the contact-less technique optical coherence elastography (OCE) has been enhanced to achieve spatial resolution of elastic properties at the scale of 20–40 µm, which achieves the range of single-cell sizes but prohibits subcellular resolution.^28,29^ Indirect techniques such as fluorescence recovery after photobleaching (FRAP) relate the diffusion coefficient to mechanical properties but compromise spatial resolution and sensitivity.^30^ Brillouin microscopy measures inelastic scattering of an incident laser by its interactions with spontaneous, thermally induced acoustic phonons (propagating density fluctuations) that depend on elastic and viscous properties of the assessed material.^31,32^ Being label-free and non-invasive, it permits repeated measurements of macroscopic volumes while resolving features at and below cell size and is therefore suited to follow the temporal evolution of imprinted patterns of heterogeneity.^31^

This work introduces Mechanoscribed Xolography, which combines grayscaled dual-color VAM with in-situ object localization, to prescribe local differences in crosslink density directly into cell-laden hydrogels during printing. These differences are intrinsic to the printed matrix, so that embedded cells encounter them without the need for external loading. The advance over prior work is threefold: First, grayscaling is achieved in a cytocompatible, cell-laden formulation with imprinted heterogeneity being maintained despite post-curing and for a culture period of three weeks, which has not been demonstrated before. Second, printing and grayscaling are performed relative to detected objects during printing, without the need for fluorescent labeling, so that the environment of a cellular spheroid can be modified rather than the construct as a whole. Third, the imprinted pattern is verified non-destructively over the culture period with Brillouin microscopy, which resolves the evolution of material properties and cell–material interactions within a single construct over time. Hydrogels with regionally distinct crosslinking regions produce correspondingly distinct cell behavior in terms of cell morphology and differentiation behavior (Figure 1). This work provides a novel pipeline for high-precision fabrication and assessment of cell-laden constructs with spatially controlled mechanical properties that drive heterogeneous cellular response and thereby opens unique opportunities for mechanobiology research and engineering of biomaterials or tissue models for tissue engineering and regenerative medicine.

**Figure 1.**
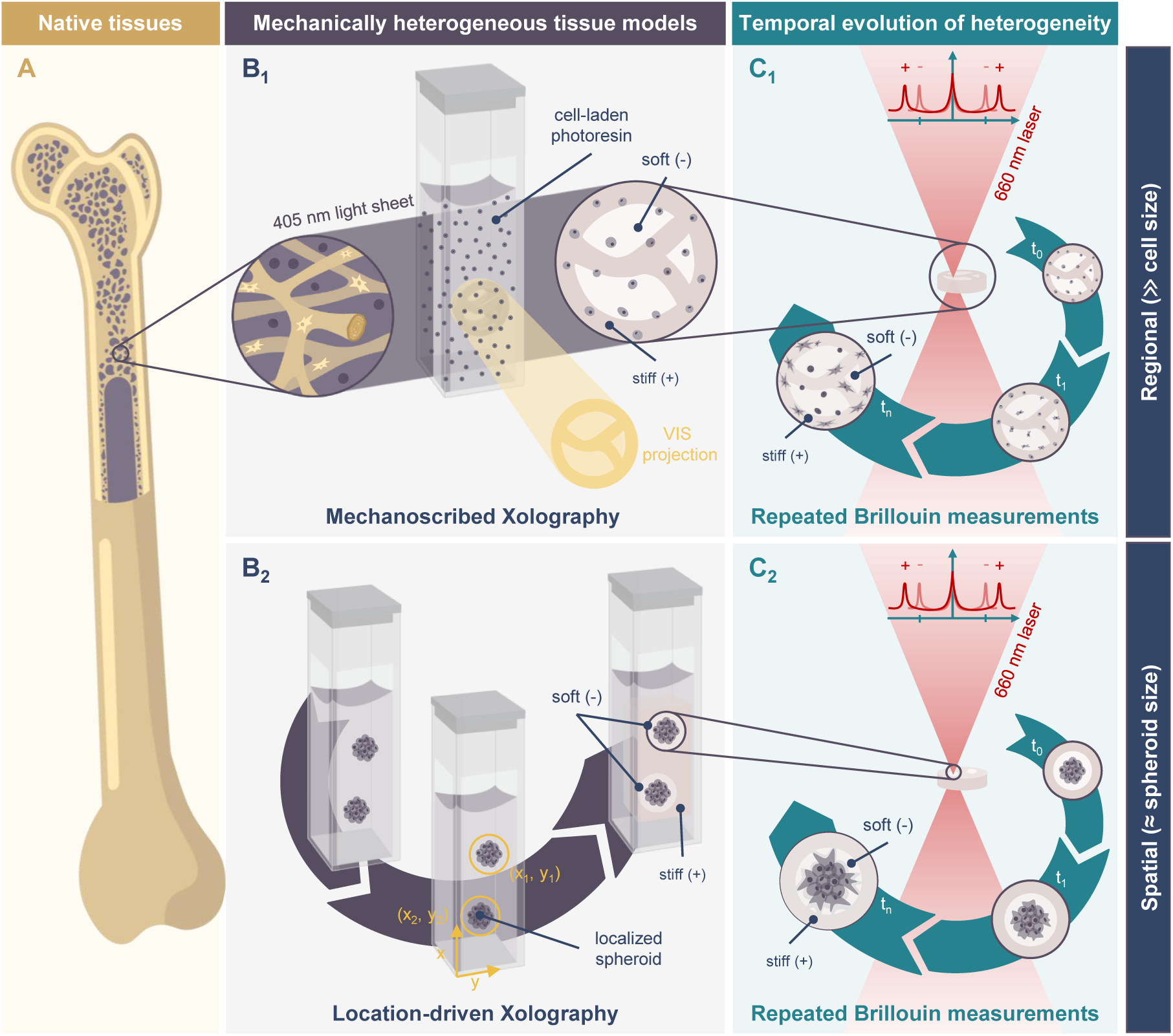
Schematic workflow of this study: We introduce Mechanoscribed Xolography to manufacture cell-laden hydrogels with controlled mechanical heterogeneity and combine this approach with Brillouin microscopy to gain regional and local control of volumetric mechanical properties and track their spatiotemporal development. A) Inspired by mechanical heterogeneity e.g. present between trabeculae and the surrounding bone marrow, B_1_) Mechanoscribed Xolography was used to create mechanically distinct regions, stiff (+) and soft (-), in 3D hydrogel models. C_1_) Properties of cells and hydrogels were monitored by live Brillouin measurements over time and guidance of cellular behavior was confirmed with fluorescent microscopy at culture end points. The results demonstrated varying cell behavior between non-grayscaled and grayscaled regions. B_2_) A localization-driven printing pipeline was implemented to allow the detection of objects like cellular spheroids within the photoresin and enable the selective mechanical modification of their proximate environment. C_2_) Growth of selected spheroids was tracked over time with repeated Brillouin microscopy.

## 2. Results and discussion

### 2.1. Achieving heterogeneous material properties in cytocompatible 3D hydrogels

Building on recent advances in cell-laden photoresin formulations for Xolography, this work comprised a GelMA-based hydrogel with BIS-TRIS as the coinitiator, a transparent and commercially available Dulbecco’s Modified Eagle’s Medium (DMEM) formulation specifically developed for fluorescent imaging, and iodixanol as a refractive index matching agent. Together, the components are referred to as the base formulation. In a second formulation, referred to as the Biobooster formulation, TEMPO-OH, a radical scavenger and a crosslinking enhancer called Biobooster, was added to the base formulation (Figure 2 A). Given the negligible impact of omitting Fetal Bovine Serum (FBS) during printing on both viability and metabolic activity of hMSCs, as well as its high absorbance below wavelengths of 425 nm, the photoresin herein was intentionally not supplemented with FBS.^22^ The refractive index of the cell-free photoresin after addition of iodixanol was measured to be 1.38 and thus matched the refractive index of cytoplasm in the range of 1.37–1.38.^33^ In contrast to previous work which utilized an ultraviolet light sheet of 375 nm wavelength, a violet light sheet of 405 nm wavelength was introduced for bioprinting in this work (Figure 2 B).^20–22^ This switch supported the selective activation of DCPI without compromising on penetration depth or causing light attenuation due to iodixanol addition (Figure S1 A). Besides the improved optical performance and the general reduction in scattering, the major benefit of the violet compared to the ultraviolet light sheet previously utilized for Xolography lies in potentially decreased DNA damage.^34^

**Figure 2.**
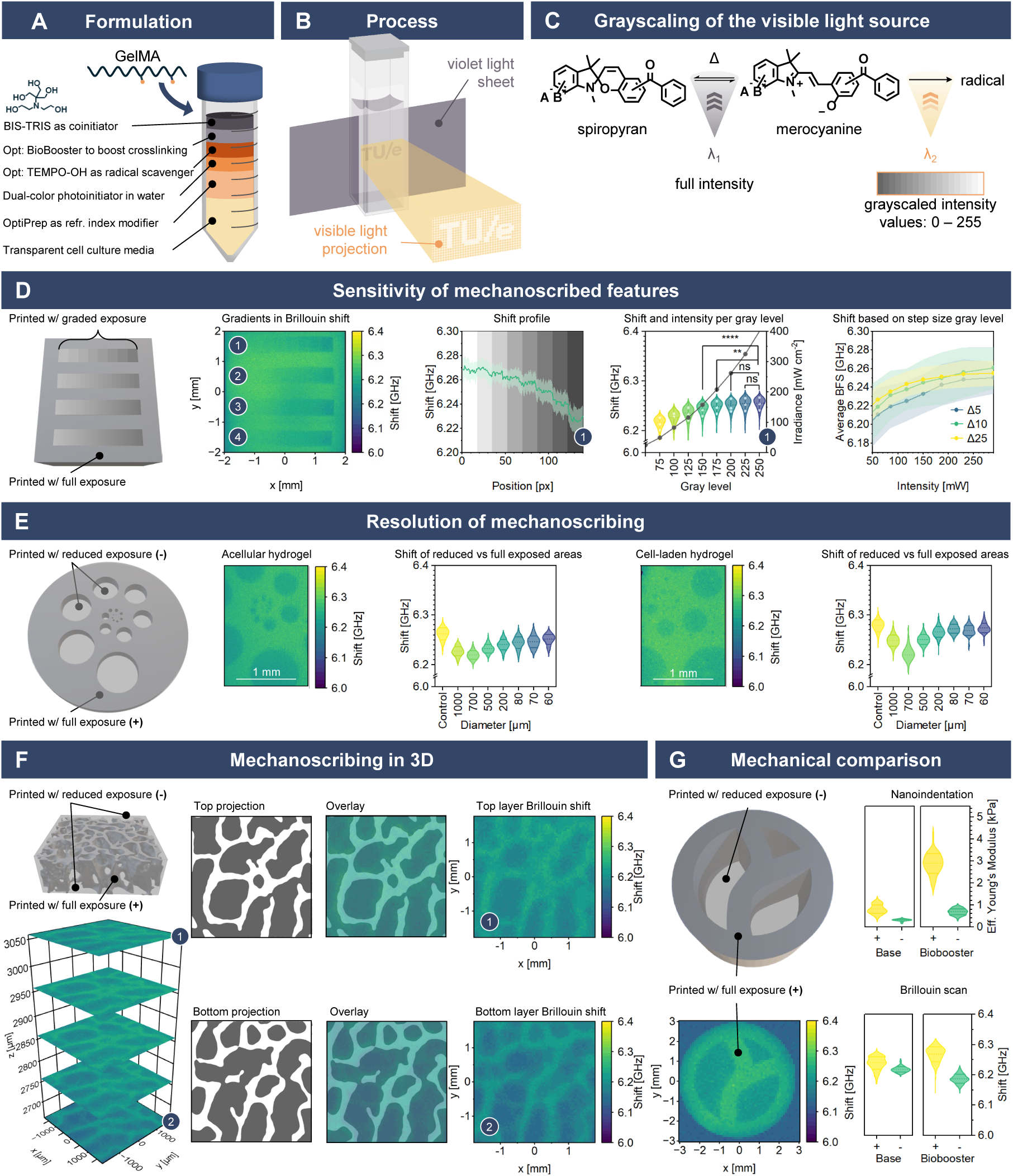
Three-dimensional control of mechanical properties with a cross-sectional resolution of 70 µm in cell-free and 200 µm in cell-laden hydrogel models through Mechanoscribed Xolography was suggested by a multi-modal characterization approach with Brillouin microscopy and nanoindentation. A) The utilized base-formulation photoresin was composed of GelMA, the photoinitiator, the coinitiator BIS-TRIS, iodixanol solution, and transparent culture media. Additionally, a Biobooster formulation also included TEMPO-OH and diaryliodonium salt. B) The principle of Xolography is based on dual-color photopolymerization, herein initiated by the orthogonal intersection of a violet light sheet and a visible light projection. C) The initially dormant photoinitiator (spiropyran isomer) gets excited by irradiation with 405 nm light and transitions to its latent, activated form (merocyanine isomer), in which incident visible light in the presence of a coinitiator triggers radical formation and photopolymerization. Grayscaling of the visible light results in spatially defined light-dosing which in turn affects material properties like stiffness. D) Grayscaling resulted in non-linear modulation of material properties as measured by BFS shift. The BFS profile shows a representative hydrogel (*N* = 1); BFS per gray level is shown in a violin plot, with each point representing one hydrogel (*N* = 3) E) BFS was dependent on the dimensions of grayscaled features and the contrast between a grayscaled region and fully exposed surroundings decreases with decreasing feature size. For a gray level of 100, differences were maintained down to a cross-section of 70 µm and 200 µm in acellular and cell-laden hydrogels, respectively. n = 18822, 36, 1617, 1109, 257, 55, 28, 19 (acellular) and n = 18221, 3868, 1617, 1405, 266, 61, 32, 27 (cell-laden) pixels from N = 1 hydrogel. F) Mechanical properties were tunable in 3D, as seen in the printed sample showing an extract of a trabecular bone scan obtained from a human proximal femur (CAD geometry licensed under CC BY 4.0), which was printed at actual trabecular feature sizes. Dimensions of the bone sample were 6 mm *×* 6 mm *×* 2 mm, representative BFS maps show patches of 4 mm *×* 4 mm of the print, sampled with a height difference of 0.4 mm. N = 1 hydrogel. G) Lower BFS shifts (n = 556, 141 (base), 78 (Biobooster) measured pixels from N = 3 (base), 3 (Biobooster) hydrogels) for grayscaled (-) areas compared to non-grayscaled (+) areas corresponded to lower Young’s moduli as mapped by nanoindentation (n = 30, 60 indentation curves from N = 1 (base), 3 (Biobooster) hydrogels). Data shown in violin plots as mean *±* SD, statistical analysis was performed with a Kruskal–Wallis test with post-hoc Dunn’s pairwise comparison, p > 0.05 (ns), p < 0.05 (*), p < 0.01 (**), p < 0.001 (***), p < 0.0001 (****).

The underlying principle of grayscaled Xolography along the dual wavelength-initiated photopolymerization scheme is visualized in Figure 2 C. The DCPI remains mainly unresponsive to visible (VIS) light in its dormant form (spiropyran isomer). Incident violet light (*λ*_1_ = 405 nm) induces photoisomerization of spiropyran to the unstable active merocyanine isomer. In its merocyanine form, the DCPI is excited by VIS light (*λ*_2_, Figure S1 B) and initiates radical photopolymerization in the presence of a co-initiator serving as an electron–donor. The violet light is applied as a light sheet, enabling in-plane activation of the DCPI, while photopolymerization is induced via an orthogonally projected VIS-pattern. This configuration spatially confines polymer crosslinking to the intersecting region of both wavelengths (Figure 2 B). The printing speed, which corresponds to the exposure time, and the intensity of the violet light sheet determine the available concentration of merocyanine species. The printing speed and intensity of the intersecting VIS-projection in turn affect radical formation and, respectively, local polymerization. The VIS-projection represents successive slices of the CAD model at every position and distinguishes in traditional binary Xolography pixel-wise between full irradiation within the printed object (white pixels) and no irradiation surrounding the object (black pixels). In contrast to this, grayscaled Xolography uses an 8-bit spectrum (black = 0, gray = x with x ∈ {1,254}, white = 255) which results in an adjustable intensity of irradiation between full and zero exposure (Figure S2). Notably, the measured irradiation intensity scaled non-linearly with gray level, following a power law, *I* ∝ *g^γ^* with *γ >* 1, where *g* is the gray level imposed by the projector’s gamma encoding.

To balance construct stability during cell culture with the maintenance of imprinted mechanical heterogeneity, we investigated post-curing after overnight washing as an alternative to intraday post-curing (Figure S3). Samples that were post-cured after overnight incubation in PBS or cell culture medium preserved a wide range of mechanical properties between regions of full and reduced exposure. At the same time, mechanical properties were overall slightly increased, which was a necessary requirement for longer hMSC culture: Grayscaled areas of non-postcured but cell-laden parts were completely degraded after 9 days in culture (Figure S4), likely due to secretion of matrix proteinases (MMPs) that enzymatically digested the GelMA network.^35–37^ Although areas that were crosslinked with full exposure to the VIS-light source were also affected during cell culture, they demonstrated integrity at day 9. Therefore, all subsequent prints were post-cured after overnight washing.

To investigate the impact of grayscaling in Xolography on spatial and temporal material properties, Brillouin microscopy was utilized. The frequency shift of the detected Brillouin peaks (BFS) in Brillouin microscopy is a mean to calculate the material longitudinal storage modulus (M’), and the width of the peaks is related to the longitudinal loss modulus (M”). However, M’ and M” depend on the refractive index (n) and the density (*ρ*) of the material, which is challenging to obtain in heterogeneous samples like biological specimens.^31^ Simplifying assumptions that tried to establish a relationship between the refractive index and the density of a biological sample, such as *n*^2^ ∝ *ρ*, were falsified for samples with lipid-containing adipocytes.^38–40^ Furthermore, there is no universal relationship between the longitudinal modulus M probed by high frequencies and the Young’s modulus E, but empirical, context-dependent relationships were found.^31,41^ Therefore, BFS was used herein as a proxy for M’ to compare material properties within a single sample over time rather than as an absolute value for elasticity.

By evaluating the spatial BFS of different sample geometries, we initially determined the boundary conditions of mechanical control in the utilized GelMA formulation. A test structure comprising four gradients with decreasing step sizes between gray levels (25, 10, 5, and quasi-continuous), spanning values from 80 to 250, was designed to assess the sensitivity of mechanical control (Figure 2 D). Gray levels below 50 were excluded due to insufficient crosslinking (Figure S5). An analysis of the BFS corresponding to gradient N*^o^* 1 with the largest investigated step size between the gray levels indicated that no significant differences were measured for gray levels between 200 and 250, while all other levels were significantly different. The asymptotic behavior with increasing gray levels pointed towards a saturation of the degree of crosslinking achievable in this combination of materials and processing settings. To balance printability and enlarged mechanical contrast, a lower gray level of 100 (20 % of full irradiance) was selected for subsequent experiments. The gradient with the smaller step size of 5 exhibited a slightly irregular trend, while gradients with larger step size (10, 25) showed an inverse dependence on intensity, which can be described by the frequently observed reaction kinetics of radical photopolymerization. The signal-to-noise ratio for the smaller step-sized gradient might have suppressed this effect but could also be an artifact of process-inherent in-plane inhomogeneities in the degree of crosslinking (DoC).^42^

We further investigated the resolution of mechanical control in the projection plane. Although theoretically, local grayscaling could be implemented at the projector pixel size of 4.6 µm, the actual resolution of grayscaling is limited by light scattering and diffusion processes during photopolymerization. To identify these effects in the GelMA-based photoresin, we determined the minimum required diameter of grayscaled cylinders in a printed disk that still resulted in a measurable difference in mechanical properties (Figure 2 E). In a cell-free GelMA hydrogel, the BFS measured in cylinders with diameters down to 70 µm appear different compared to the surrounding non-grayscaled hydrogel as the control. The contrast in BFS diminished with decreasing diameter, likely due to radical diffusion across boundaries. Similar behavior was observed in a cell-laden sample, however, scattering reduced BFS contrast at smaller scales and the shift for grayscaled features of 80 µm diameter and below was shown to be similar to the shift of the surrounding, fully exposed hydrogel.

To demonstrate the potential of grayscaled Xolography to manufacture complex 3D structures with heterogeneous properties, a trabecular bone structure was printed at actual size and evaluated (Figure 2 F, Video S1). Grayscaling improved fabrication integrity: softer inter-trabecular regions provided structural support during washing and post-curing, preventing the collapse observed in non-grayscaled designs with void spaces between trabeculae (Figure S6 C). This resulted in a notable improvement in feature representation.^43^ Notably, the BFS map closely matched the projected grayscale patterns across the measured construct height. Brillouin microscopy correctly identified 96 % of the stiff (bone) voxels and 71 % of the soft voxels, with 80 % of all voxels classified correctly (Figure S6 D, E). Considering the different degradation profiles of grayscaled and non-grayscaled regions before post-curing (Figure S4), this strategy opens unique possibilities to aid manufacturability while enabling the generation of void spaces through sacrificial, grayscaled regions for perfusion or vascularization in cellular constructs.

Lastly, to ensure that the previously obtained BFS shifts were in accordance with regions of different mechanical properties, nanoindentation was performed on a simplified trabecular-like structure (Figure 2 G) containing grayscaled (gray level 100) and fully exposed (gray level 255) regions. The effective Young’s moduli were 313 ± 46 Pa and 814 ± 200 Pa (identical loading rate; absolute values are rate-dependent given GelMA’s viscoelasticity) for the grayscaled and fully exposed regions printed with the base formulation, respectively, confirming that grayscaled regions were significantly softer than the fully exposed regions. To increase the crosslinking density and thus extend the accessible range of mechanical properties, Biobooster, a water-soluble and cytocompatible diaryliodonium salt, was added to the formulation. In three-component photoinitiating systems, diaryliodonium salts act as electron acceptors that oxidize the reduced photoinitiator, regenerating it and generating additional initiating radicals.^21^ Here, the system is composed of DCPI, BisTris, and the Biobooster. The resulting increase in reactivity is partly compensated by the addition of radical scavenger TEMPO-OH to maintain print resolution.^44^ Hydrogels printed with the Biobooster formulation resulted in an increased effective Young’s moduli of 690 ± 137 Pa and 2891 ± 454 Pa for the grayscaled and fully exposed regions. Correspondingly, Brillouin microscopy showed significantly lower BFS in the grayscaled regions for both formulations. The elasticity range covered here corresponds roughly to the elasticity of the central bone marrow (0.3 kPa) and endothelia (0.5–2 kPa), and was thereby severely softer than the typical stiffness of bone in the range of GPa.^45,46^ However, bone formation is known to start from relatively soft (25–40 kPa) osteoid matrix composed of crosslinked collagen and eventually stiffens over weeks through mineralization into bone tissue.^45,47,48^ Furthermore, bone fracture healing is marked by hematoma formation with an elasticity of about 50–200 kPa within the initial 14 days.^48^ Notably, despite the addition of Biobooster, the stiffness range obtained herein was below the known stiffness range at which MSCs preferably differentiate into osteogenic lineage in culture (> 20 kPa).^11,47^ To assess whether gray level also affects network properties beyond stiffness, sol fraction and mass swelling ratio were determined for samples printed with the base formulation (Table S1). After post-curing, gray levels 100 and 255 showed comparable sol fractions (37 % and 39 %) and mass swelling ratios (approximately 15). Moreover, in contrast to the Brillouin shift, maps of the Brillouin Full Width Half Max (FWHM), a proxy for the viscous component, showed no clear pattern (Figure S7). These results suggest that, within the resolution of these measurements, the gray levels here differed mainly in stiffness. Nevertheless, other network parameters, such as mesh size and degradability, are inherently linked to crosslink density and cannot be varied independently in the present material system.

### 2.2. Material properties evolve during three weeks of culture but maintain imprinted differences

We performed repeated Brillouin scans of the same hydrogels with embedded hMSCs during a 21-day culture period to study the evolution of material properties in stiff and soft regions in different media conditions, i.e. different biochemical stimulation (Figure 3 A–P). The chosen media compositions were known to drive hMSC differentiation to different degrees: an osteogenic (O) and adipogenic (A) medium were supposed to predominantly drive hMSC differentiation towards osteogenic and adipogenic lineages, respectively.^11^ A full (F) media composition containing both osteogenic and adipogenic cues was intended to biochemically equilibrate osteogenic and adipogenic differentiation and transition hMSCs to a state in which mechanical cues imposed by grayscaling sufficed to determine differentiation fate. A maintenance medium (C) for hMSCs without growth factor supplementation was selected as a control. Additionally, acellular hydrogels cultured in maintenance medium were included as a second control. The printed samples contained three zones exposed to lower visible light intensity (gray level 100), which remained distinguishable by visual inspection across all media conditions three days after printing. Over time, a decrease in BFS was observed across all media conditions (Figure 3 Summary, Video S2), which was significant for acellular, adipogenic and osteogenic constructs (Two-Way ANOVA, time) and may reflect matrix degradation as well as changes in local refractive index and density. Since the squared refractive index scales approximately with mass density, their variations largely cancel in BFS, except in lipid-rich or mineralized regions, as may arise under adipogenic and osteogenic conditions.^31,38^ Therefore, we do not interpret the BFS of cell-laden constructs biologically. Acellular constructs showed a comparable decrease in BFS, as the temporal change did not differ significantly between acellular and the cell-laden constructs (Three-Way ANOVA, time × media interaction), and nanoindentation confirmed a decrease in Young’s modulus between day 4 and 21 (Figure S8). This indicates that the temporal decrease is predominantly caused by intrinsic changes of the hydrogel network rather than by cell-driven remodeling.

**Figure 3.**
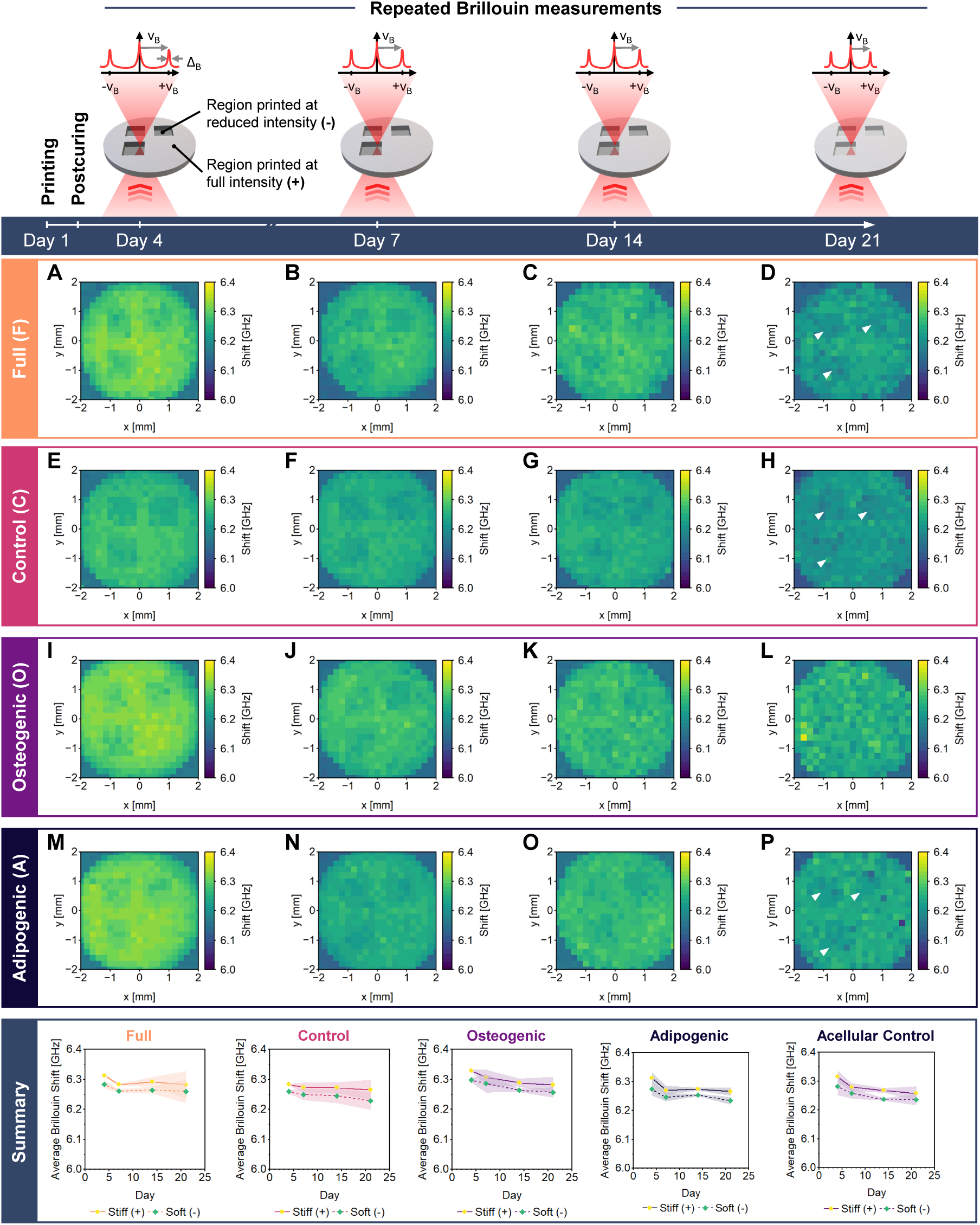
Repeated Brillouin measurements of the same scaffold illustrated the development of material properties in different culture conditions over time. Although the BFS decreased consistently regardless of culture supplementation, a measurable difference between the grayscaled regions and their surroundings persisted. The top row schematically indicates the measurements on day 4, 7, 14, and 21 after printing and the general development of BFS and the scaffold’s visual appearance over time. A–D) Temporal changes of the BFS map of a representative single grayscaled hydrogel scaffold cultured in medium containing full supplementation to induce biochemical balance between osteogenic and adipogenic differentiation. In comparison, temporal changes of the BFS map of a scaffold cultured in standard hMSC maintenance medium (E–H), in osteogenic-supplemented media (I–L), and in adipogenic-supplemented media (M–P). Summary) The average BFS for fully irradiated, stiff (+) areas and grayscaled, soft (-) areas printed at lower visible light intensity (gray level 100) per culture condition showed a general decrease in BFS over time while maintaining the difference between grayscaled and non-grayscaled areas. One measurement of the stiff/soft interface of each square was taken. Data shown as mean *±* SD in error bands, N = 3 hydrogels from 2 independent experiments.

Although the visual differences between stiff and soft regions diminished over time, measurable differences in material properties were maintained. In all pairwise comparisons with acellular constructs, the stiff regions showed a significantly higher BFS than soft regions (Three-Way ANOVA, region), and this difference did not change significantly over time (time × region interaction) or between the acellular and cell-laden constructs (media × region interaction). Of the cell-laden conditions, only osteogenic constructs differed significantly from the acellular condition. Within individual conditions, the difference between regions was significant for acellular and adipogenic constructs (Two-Way ANOVA, region), but did not show significance for full, osteogenic, and control constructs, likely due to the higher variability between samples. Overall, the hydrogels exhibited time-dependent changes in material properties, largely independent of the presence of cells and the media condition while maintaining locally imprinted BFS differences up to 21 days in culture. Based on these findings, we proceeded to further investigate the impact of grayscaled regions on cellular response.

### 2.3. Mechanoscribed Xolography impacts cellular response

To investigate whether grayscaled regions (effective Young’s modulus 313 ± 46 Pa and 814 ± 200 Pa) influence cell response, we analyzed the morphology of hMSCs 3 and 13 days after printing with our base formulation (Figure 4 A, B). Cells on the hydrogel surface exhibited the characteristic spindle-shaped morphology typical of hMSCs cultured on standard cell culture plastic. After 3 days, hMSCs on the surface of stiff regions showed a larger median area of 475.5 µm^2^ as opposed to 390.8 µm^2^ on soft regions, a trend that became more pronounced after 13 days. Slightly varying distributions of solidity and eccentricity converged during culture time. The combination of all investigated parameters suggested that cells on stiffer surfaces tend to spread slightly slower but grow bigger. Cells cultured for 21 days with samples printed with the Biobooster formulation (Figure 4 K-N) tended to exhibit a larger spread-out morphology on the soft regions at the surface, while a smaller, partially spindle-shaped morphology was seen on the stiff regions. The results of the soft area here are consistent with the base formulation samples, as nanoindentation measurements (Figure 2 G) of the stiff areas of the base formulation showed a similar effective Young’s modulus as for the soft areas of the Biobooster formulation.

**Figure 4.**
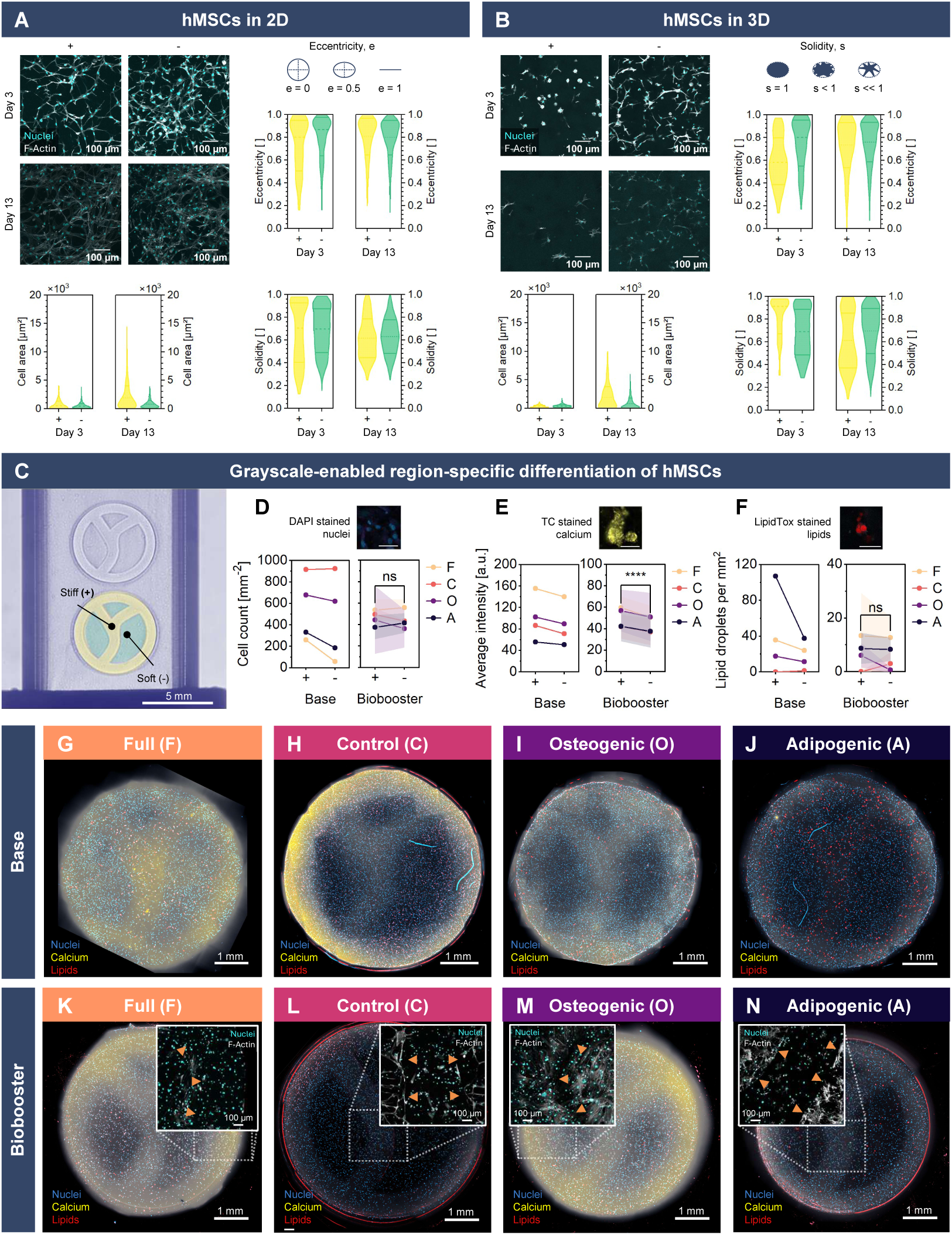
Modulated material properties through Mechanoscribed Xolography resulted in increased solidity and reduced fate decisions of hMSCs in grayscaled areas, independent of biochemical supplementation. Confocal images showing nuclei (DAPI, blue) and actin cytoskeleton (white) of hMSCs, with images acquired A) on the surface (2D) and B) at a depth of 100 µm in the hydrogel (3D) as well as quantitative assessment of the morphological features cell area, eccentricity, and solidity. (+) corresponds to stiff and (-) to soft regions in all images. A) n = 280, 602, 375, 888 cells from 3 images taken at different locations in N = 1 hydrogel. B) n = 187, 332, 110, 619 cells from 3 images taken at different locations in N = 1 hydrogel. Data shown in violin plots as mean *±* SD. C) Schlieren image of grayscaled samples with stiff and soft regions in the cuvette after printing: the simplified trabecular-like structure introduced in Figure 2 G was used to assess effects on differentiation behavior within one sample. D) Cell count per region (DAPI) was slightly higher in grayscaled regions for the base formulation, except in control medium, and did not differ significantly for the Biobooster formulation. E) Calcium deposition (TC, yellow) was higher in media with osteogenic cues and in stiff areas for all media (significant for the Biobooster formulation). F) Lipid droplets per mm^2^ (LipidTox, red) were most abundant with adipogenic cues and higher in stiff areas for the base formulation only. Scale bars 50 µm; mean *±* SD, scaffold rim excluded; *N* = 1 (base), *N* = 3 (Biobooster). Biobooster: Two-Way ANOVA with repeated measures (media *×* region); **** *p <* 0.0001 (region effect); ns, not significant. G–J) Immunofluorescent images from one side of the scaffolds printed with the base formulation cultured in G) full medium containing osteogenic and adipogenic supplementation, H) control culture medium without differentiation-related supplementation, I) cultured media that contained osteogenic supplementation, and J) media with adipogenic supplementation. K–J) Immunofluorescent images from one side of the scaffolds printed with the Biobooster formulation cultured in the same media types as the scaffolds in G–J. Overlayed images show confocal images of hMSC nuclei and cytoskeleton at the hydrogel surface at the interface between soft and stiff areas, indicated by the orange arrows.

A more visual change in morphology was observed for encapsulated cells (Figure 4 B): After one week of culture (base formulation), hMSCs in soft regions spread into spindle-shaped morphologies, whereas hMSCs in stiff regions largely maintained a circular shape. This difference in eccentricity diminished after two weeks in culture, a time point at which cells in both areas showed similar spindle-shaped morphologies. Lower cell solidity in stiffer hydrogel regions indicated the formation of more pronounced protrusions. The larger projected area and higher eccentricity observed in the stiffer zones may indicate a bias towards osteogenic differentiation.^49,50^ Likewise, low solidity in stiff zones may reflect cytoskeletal remodeling associated with osteogenic differentiation, migration, or mechanically active cells.^51^ On the contrary, higher solidity and smaller areas have been linked to adipogenic differentiation or maintenance of stemness.^52^

To further assess the influence of grayscaling on cell differentiation, we investigated hMSC behavior in the simplified trabecular-like structure (Figure 4 C) previously introduced (Figure 2 G). Analysis focused on the inner circular and Y-shaped regions. Scaffolds printed with the base and Biobooster formulation were stained for calcium deposition, an indicator of osteogenic differentiation, and lipids, an indicator of adipogenic differentiation.^53^ Representative images obtained from fluorescent widefield microscopy of the scaffolds cultured in different media conditions are provided in Figure 4 G–J. We quantified cell count, calcium deposition (based on fluorescent intensity), and the number of lipid droplets normalized by area on both sides of the scaffolds (Figure 4 D–F). Hydrogels printed with the base formulation cultured in the hMSC maintenance medium and the osteogenic medium exhibited large variations in cell count on both sides of the scaffold. Across all conditions, a higher cell number was observed in the soft regions. In contrast, cell count for the Biobooster formulation was consistent between samples and did not differ significantly between stiff and soft areas. Live/dead staining of the Biobooster samples showed predominantly viable cells in both stiff and soft regions (Figure S9). Calcium deposition for both formulations was found to be higher in constructs cultured with osteogenic cues (both the pure and the full medium) and slightly higher in stiff areas, regardless of media supplementation. For the Biobooster formulation, the difference between stiff and soft areas was significant; for the base formulation, it indicated a trend. Lipid droplets were most abundant in hydrogels cultured in medium with adipogenic cues for both hydrogel formulations. In contrast to previous assumptions, a higher number of lipid droplets was observed in stiff regions for the base formulation samples. A similar trend was observed in samples cultured with full media supplementation, even though the total lipid counts remained below those of the purely adipogenic-supplemented samples. However, these effects were less distinct in the Biobooster samples, where the lipid count was slightly higher in the full and adipogenic media samples for both stiff and soft areas, with no significant difference between the regions. Overall, biochemical supplementation appears to dominate the differentiation behavior of hMSCs. Nevertheless, subtle but consistent trends between stiff and soft regions indicated a general increase in all differentiation markers in stiff areas, which was significant for calcium deposition in the Biobooster samples, notably despite slightly lower cell numbers in these regions in the base formulation samples. It is important to note that the mechanical differences were relatively small in the base formulation (313 ± 46 Pa and 814 ± 200 Pa), i.e. approximately one order of magnitude lower than the 2.5 to 12 kPa range reported to elicit a pronounced stiffness-dependent effect.^11^ However, the herein observed trends in morphology, calcium deposition, and lipid formation collectively support the proof of concept that Mechanoscribed Xolography enables spatial modulation of cellular responses through crosslink-density-dependent differences in the local hydrogel microenvironment and the resulting stiffness differences. Mechanoscribed Xolography could thus become a tool to mimic subtle local differences in the matrix microenvironment, as observed during morphogenesis in vivo, in contrast to in vitro models, which often exaggerate the native situation.^54,55^

### 2.4. In-situ microscopy enables location-driven Xolography

In the previous experiments, we predefined regions with mechanical properties, demonstrating their influence on cell behavior at the construct level, yet all cells within a region experienced the same mechanical environment. Predefined regions cannot account for the inherent stochasticity of biological object placement in volumetric printing. This required adapting the geometry to the object distribution rather than imposing a predefined geometric and mechanical environment. This was achieved by introducing in-situ microscopy to register the position of individual biological objects, here spheroids, enabling location-driven volumetric printing with mechanical control at the level of each detected object (Figure 5). We developed a custom imaging system, integrated into the Xolography system (Figure S10), and implemented a localization technique based on object sharpness. The implementation and localization pipeline steps are shown in Figure 5 A–B. Existing in-situ feature detection has so far required fluorescent labeling.^27^ We instead used brightfield illumination from the Xolography printer’s own projector (Figure 5 C), which keeps the method label-free. At its core, the workflow acquires a z-stack across the print volume by translating the sample along the z-axis, then localizes each object in 3D based on object sharpness. Integrating in-situ imaging with volumetric printing posed four coupled challenges: (1) spatial registration of the imaging and projection systems with the print volume, (2) a field of view spanning the full printable area (17.7 × 10 mm) while resolving objects at single-cell scale, (3) depth-dependent focal shift caused by the hydrogel–air refractive index mismatch, and (4) optical aberrations arising from operating at the limits of that field of view. We addressed registration by autonomously aligning the projector and imaging system to a common coordinate system with autofocus in the z-axis and ChArUco boards in the xy-plane (Figure 5 D, Video S3). To cover the full printable area while retaining single-cell resolution, we paired a low-magnification, long-working-distance objective with a shorter-focal-length tube lens and a large-format, small-pixel sensor. Because of the hydrogel–air refractive index mismatch, the actual focal plane shifts with depth during axial scanning. We corrected this by repositioning the imaging unit with a calculated offset, following the correction established in DescSPIM.^56^ Operating at the field-of-view limit of the objective introduced field curvature and vignetting. Optical simulation, experimentally validated, showed that off-axis objects focus progressively farther from the imaging plane; we fitted a paraboloid to the measured focal-plane positions and applied it as a per-particle z-correction (Figure 5D). Vignetting was corrected by reshaping the projector illumination profile to a uniform target (Figure S11). Object detection was validated using synthetic microparticles and cell-derived spheroids (hPACs and MSCs) across a broad size range (20–500 µm). Particles were suspended in GelMA, and both localization and printing were performed after GelMA physically gelled, which maintained spheroid position and prevented sedimentation throughout localization and subsequent printing.^19^ Polyethylene microparticles (Figure 5 E) and polystyrene microparticles spanning 500 µm to 106 µm (Figure 5 F) with 97 %–100 % precision and 74 %–97 % recall (Table S2). As a proof of concept, microparticles down to 20 µm were also successfully detected (Figure S12 A).

**Figure 5.**
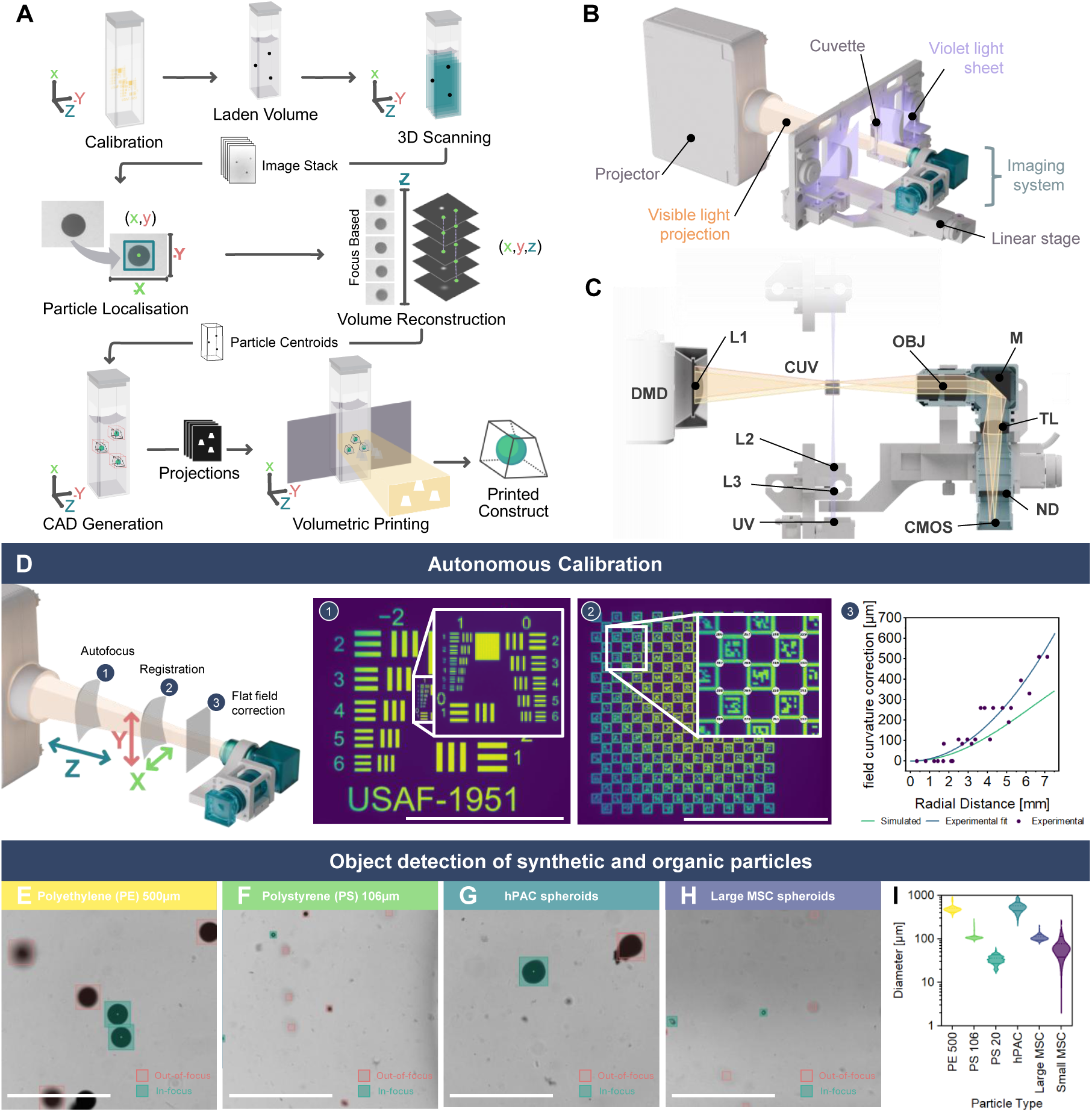
Working principle and hardware implementation of location-driven Xolography. A) Schematic overview of the particle localization pipeline, from volumetric acquisition to CAD generation and printing. B) 3D render of Xube^2^ components of projector and light sheet, including the imaging system. C) Top-down detailed 3D render of the imaging system integrated within the Xube^2^, with the optical path indicated schematically relative to the printing optics. D) Autonomous calibration steps consisting of (1) Z-alignment via projected USAF 1951 target; (2) XY-alignment of projector–imaging unit, through ChArUco board projections; (3) Flat-field correction using experimentally found field curvature compared to Zemax simulated field curvature. Scale bars are 5 mm. E–H) Representative images for object detection overlaid on cuvette as observed by imaging samples of 500 µm polyethylene particles (E), 106 µm polystyrene particles (F), hPAC spheroids (G) and large MSC spheroids (H). Scale bars are 2.5 mm. I) Localization-detected particle diameter distributions per population, number of total particles found: 23, 170, 8286, 8, 167, 36, respectively (violin plot, log-scaled y-axis). N *≥* 4 independent samples, except for Small MSC (N = 2) and PS 20 (N = 1)

For biologically relevant objects, human primary articular chondrocyte (hPAC) spheroids were produced and imaged as a large-spheroid model (Figure 5 G) and MSC spheroids as a smaller spheroid model (Figure 5 H) with 97 %– 100 % precision and 64 %–65 % recall (Table S2). Smaller Y201 MSC spheroids of approximately 50 µm were also resolved (Figure S12 B).^57^ Segmentation-derived diameters were used to confirm that detected objects corresponded to the intended size population rather than smaller or larger artifacts. A slight overestimation was expected from thresholding at the mask boundary, but the measured diameters of each population matched its known size, confirming correct detection (Figure 5 I). The typical time required for the localization pipeline up to the point of printing is reported in Table S3. Benchmarking up to ∼ 10^6^ particles mL*^−^*^1^ (Figure S13) showed that frame-to-frame particle linking scales least favorably with particle density. This was not limiting at the densities used in this work. For higher densities, such as those relevant to single cells, linking can be replaced by 3D proximity clustering of the detections. Together, these results demonstrate label-free localization of synthetic and biological objects across the biologically relevant range from single cells (∼ 20 µm) to organoids (∼ 500 µm), characterized over 106 to 500 µm and confirmed down to 20 µm, establishing object detection within the volumetric printing process.

### 2.5. Selective modification of the mechanical microenvironment of spheroids

Localization-driven printing with mechanoscribed encapsulation was first demonstrated by encapsulating synthetic particles within Xolography-printed constructs (Figure 6 A–D, Video S4). To maintain structural integrity, the exterior shell was printed stiff, and a smaller inner trapezoid exposed each particle to a grayscaled microenvironment (Figure 6 A).

**Figure 6.**
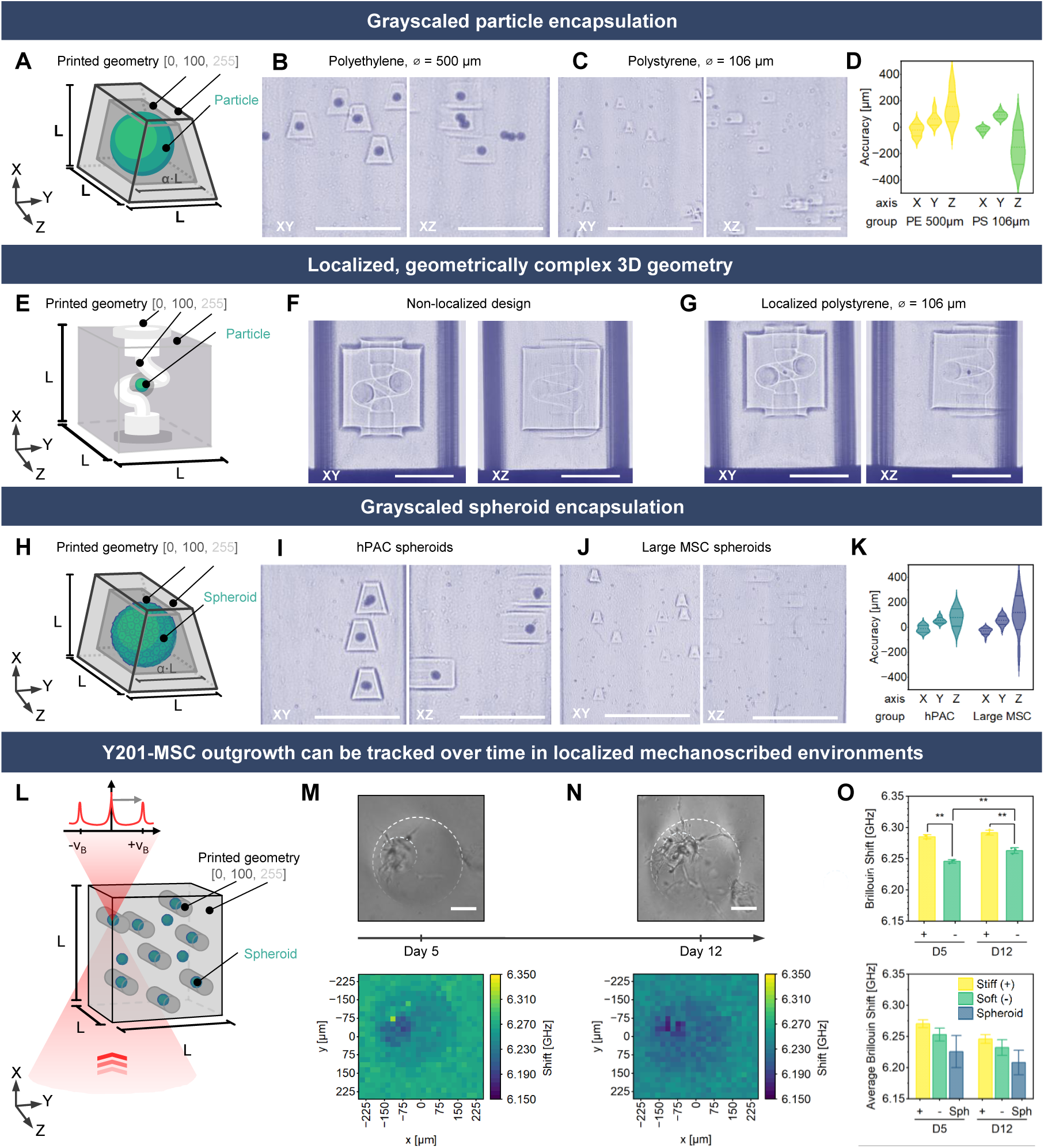
Mechanoscribed Xolography allows for the in-situ modulation of the 3D environment of randomly distributed particles within the print volume. A) Schematic of CAD design of grayscale trapezoidal encapsulation of synthetic particles B, C) Schlieren images of encapsulation of polyethylene particles (B, ∅ = 500 µm, *l* = 1 mm, *α* = 0.75), and polystyrene particles (C, ∅ = 106 µm, *l* = 750 µm, *α* = 0.75). D) Violin plots of per-axis printing accuracy, measured as the absolute centroid-to-centroid distance between each particle and its surrounding printed feature, measured from Schlieren images. N = 3 samples E) CAD model of perfusable bioreactor (*l* = 7 mm). F, G) Schlieren images of the bioreactor printed with non-localized geometry (F) and with localized encapsulation of a 106 µm polystyrene particle (G) H) Schematic of CAD design of grayscale trapezoidal encapsulation of spheroids I, J) Schlieren images of encapsulation of hPAC spheroids (I, *l* = 2 mm, *α* = 0.75) and MSC spheroids (J, *l* = 750 µm, *α* = 0.75). K) Violin plots of per-axis printing accuracy, measured as the absolute centroid-to-centroid distance between each particle and its surrounding printed feature, measured from Schlieren images. N = 3 samples L) Schematic of CAD design of encapsulated MSC spheroids. M) Brightfield image and corresponding BFS maps of an MSC spheroid’s outgrowth 5 days after printing show growth mainly in the soft environment. N) Brightfield image and corresponding BFS maps of an MSC spheroid’s outgrowth 12 days after printing show growth in both the stiff and the soft environment. O) Top: Average BFS of control scaffolds containing non-localized spheroids. Bottom: Average BFS for the grayscaled proximal zone, the non-grayscaled surrounding, and the spheroid. Scale bars are 5 mm unless stated otherwise; scale bars in panels M) and N) are 100 µm. N = 1 hydrogel (proof-of-concept); values from repeated measurements of one spheroid.

Printing across the full z-range elongated objects beyond their design (Figure S14 A), caused by out-of-plane polymerization in regions already exposed by the light sheet. We suppressed this with 4-hydroxy-TEMPO (Figure S14 B), a water-soluble TEMPO-class radical inhibitor used to improve resolution in volumetric printing.^58,59^ Its cytotoxicity was characterized separately and showed no detectable effect at the working concentration (Figure S14 C,D), supporting cell-laden use. Larger constructs required lower UV intensity to crosslink than smaller features at fixed scan speed (Figure S14 E,F); UV intensity was therefore matched to each printed part, whose size was scaled to the object it encapsulated. Schlieren imaging resolved the refractive-index contrast between partial and fully exposed regions, confirming the grayscale patterned regions (Figure 6 B, C). For both particles, their low optical transparency cast a projector shadow, leaving under-crosslinked voids on the non-illuminated side (Figure 6 B), an artifact specific to optically opaque objects. Particles aligned along the optical axis are resolved when separated by more than approximately *d*_p_ + DoF + Δ*z* (≈ 244 µm for 100 µm spheroids), making axial merging negligible at the densities used in this work, with an estimated ∼0.1 % of spheroids affected at 100 mL*^−^*^1^. Localization accuracy, taken as the centroid offset between particle and construct, was small in-plane for both synthetic materials but larger axially (Figure 6 D, Table 1).

**Table 1.** Centroid offsets between detected and printed positions for synthetic microparticles and biological spheroids. Values are mean *±* SD in µm.

| | Object | X [$\mu\text{m}$] | Y [$\mu\text{m}$] | Z [$\mu\text{m}$] |
| --- | --- | --- | --- | --- |
| Synthetic | PE, 500 $\mu\text{m}$ | $-21 \pm 44$ | $61 \pm 46$ | $150 \pm 110$ |
| | PS, 106 $\mu\text{m}$ | $-16 \pm 20$ | $90 \pm 26$ | $-150 \pm 130$ |
| Biological | hPAC spheroids | $-11 \pm 28$ | $55 \pm 23$ | $77 \pm 73$ |
| | Large MSC spheroids | $-30 \pm 24$ | $56 \pm 35$ | $120 \pm 140$ |
| Pooled | | $-21 \pm 27$ | $67 \pm 37$ | $10 \pm 175$ |

**Table 2.** Media supplementation for cell culture differentiation in printed 3D hydrogels.

| Component | Full |  | Control | Osteogenic | Adipogenic |  | Supplier |
| --- | --- | --- | --- | --- | --- | --- | --- |
|  | F | F<sub>red</sub> | C | O | A | A<sub>red</sub> |  |
| DMEM |  |  | low glucose DMEM |  |  |  | Gibco, USA |
| FBS |  |  |  | 10 % |  |  | Serana Europe GmbH, Germany |
| Penicillin/streptomycin |  |  |  | 1 % |  |  | Gibco, USA |
| β-glycerol phosphate [mM] | 10 | 10 | - | 10 | - | - | G9422; Sigma Aldrich, USA |
| dexamethasone [μM] | 1 | 0.1 | - | 1 | 1 | 0.1 | D4902; Sigma Aldrich, USA |
| ascorbic acid [μg ml<sup>-1</sup>] | 50 | 50 | - | 50 | - | - | A8960; Sigma Aldrich, USA |
| indomethacine [μM] | 50 | 50 | - | - | 50 | 50 | I7378; Sigma Aldrich, USA |
| 3-isobutyl-1-methylxanthine [μM] | 0.5 | 0.5 | - | - | 0.5 | 0.5 | I5879; Sigma Aldrich, USA |
| human recombinant insulin [μg ml<sup>-1</sup>] | 10 | 10 | - | - | 10 | 10 | I9278; Sigma Aldrich, USA |

To demonstrate compatibility with more complex architectures, we fabricated a perfusable construct integrating mechanoscribed regions around a synthetic particle, combining a soft microenvironment within a stiff structural body with open spiral channels for fluid delivery (Figure 6 E). Grayscale printing produced softer regions in-plane, with a weaker but observable contrast along Z (Figure 6 F; Figure S15 A,B). The same geometry was printed encapsulating a synthetic particle (Figure 6G), and a matched construct showed unobstructed perfusion (Figure S15 C, Video S5). Together, these establish that localization-driven printing is compatible with perfusable architectures, a step toward constructs in which spheroids experience a defined mechanical microenvironment while remaining accessible to continuous medium exchange. Because Xolography is not restricted to uniform or symmetric patterns, this workflow can be combined with adaptive channel design to route channel, in a more natural fashion, networks along multiple localized constructs, with mechanoscribing defining local mechanics within these perfusable geometries.^27^ Building on tomographic printing of perfusable organ-on-a-chip models, integrating such constructs into bioreactors could support long-term study of large or high-maintenance spheroids.^60^

Large hPAC spheroids and small hMSC spheroids were encapsulated using the inhibitor-free formulation, retained for consistency with prior biological experiments. The optically opaque hPAC spheroids cast projected shadows (Figure 6 G), whereas the hMSC spheroids produced no visible shadow in Schlieren (Figure 6 H), possibly because their small size fell below the resolution limit of the Schlieren setup. Live/dead staining confirmed maintained viability in hPAC spheroids after encapsulation (Figure S16 A–C). Biological objects showed the same accuracy and precision pattern as synthetic particles, with small lateral offsets and larger axial deviations for both hPAC and MSC spheroids (Table 1, Figure 6 K). Pooled across all encapsulated objects and calibration sessions (Table 1), X and Z showed little net bias, while the consistent positive offset in Y reflects a small residual lateral registration offset. All axes are re-aligned each session, but the lateral solution is far more reproducible, so the Z bias varies in sign between objects while the lateral offset stays consistent. This makes Z the limiting axis: the axial spread sets the smallest reliably printable environment, while the sub-40 µm lateral precision permits finer in-plane geometry.

hMSC spheroids were localized in the gel and captured in a cylindrical grayscaled region with a diameter of 300 µm (Figure 6 L). While the center of adaptive printing was notably shifted, the spheroid was captured within the grayscaled zone (Figure 6 M). The extracted BFS demonstrated differences in mechanics between grayscaled and non-grayscaled zones. After an additional week in culture, the area of the spheroid increased noticeably (Figure 6 N). As previously observed, BFS decreased over culture time but maintained the relation between shifts measured in soft areas, stiff areas, and the spheroid. Measurements of multiple interface regions in a spheroid-containing, but non-localized hydrogel are summarized at the top in Figure 6 O. Fully and partially irradiated areas demonstrated consistent differences in BFS; however, as opposed to the results of single hMSCs, the BFS increased over culture time (Figure 6 O, bottom). This trend was significant for the grayscaled area. We hypothesize it reflects the lower hMSC density in these constructs, and correspondingly reduced cell–material remodeling. Together, these prints show successful encapsulation of a spheroid within a defined mechanical microenvironment with Mechanoscribed Xolography. The defined mechanical environment and its visual distinction from the remaining, multiple millimeter-large scaffold allowed us to follow the growth of a specific spheroid over culture time with Brillouin microscopy.

## 3. Conclusion

The mechanical heterogeneity of native tissues remains insufficiently reproduced in 3D tissue models due to limited manufacturing capabilities and might severely affect experimental outcomes. This work introduced Mechanoscribed Xolography, to control elasticity in cytocompatible hydrogels globally and locally at the spheroid level (in-plane resolution of 70 µm). The contrast of imprinted heterogeneity, as quantified by BFS, was maintained over a 21-day cell culture period, despite a general decrease in BFS and across varying biochemical supplementation.

Cells embedded in heterogeneous hydrogels showed mechanically-dependent morphological changes: hMSCs in soft areas spread over a generally smaller area, while a transition from spherical to elongated cells with increased cell area was observed over ten days in stiff zones. Furthermore, differentiation behavior was enhanced in stiff zones within a hydrogel exposed to osteogenic and/or adipogenic supplementation, measured by an increase in calcium accumulation and formation of lipid droplets and associated with osteogenic and adipogenic differentiation, respectively. Notably, the approach utilized a single material system and created subtle differences in elasticity (313–814 Pa for the base and 690–2,891 Pa for the Biobooster formulation) in a range found in native morphogenesis, which in this study indicated an effect on the cellular response. The precise modulation of the mechanical environment of cellular spheroids was achieved by integrating autonomous optical microscopy into the print process. Synthetic particles as small as 20 µm were detected in the three-dimensional print volume, indicating the optical resolution of the integrated imaging system. Printing around detected objects was demonstrated down to cellular spheroids of 76 µm.

Hence, Mechanoscribed Xolography provides a rapid, high-resolution platform to create various 3D microenvironments within a single sample and enables investigation of multiple mechanical conditions within a single print, significantly reducing costs and the influence of extraneous variables on experimental outcomes. In the future, it can evolve into a tool for the experimental modeling of mechanically distinct tissues such as bone marrow or cartilage. Especially when combined with three-dimensional and non-destructive Brillouin microscopy, Mechanoscribed Xolography can be used to investigate disease progression in models of osteoporosis or fibrosis, pathological conditions associated with local mechanical changes in the extracellular matrix.

## 4. Experimental section

### 4.1. Hydrogel preparation

10 % w/v base formulation hydrogels were prepared by dissolving GelMA *(Technical Grade X-Pure GelMA 160P80 RG, Rousselot BV, Belgium)* in sterile-filtered 200 mM BIS-TRIS, 60 % w/v *OptiPrep (StemCell Technologies Canada Inc., Canada)* and DMEM *FluoroBrite (Gibco, USA)* at 45 °C. The DCPI *(DCPI 6001, xolo GmbH, Germany)* was dissolved in ultra pure water (UPW) at a concentration of 4.8 mg ml^-1^ by sonification for 20 min, sterile-filtered, and added at a final concentration of 100 ppm to the GelMA solution once both were fully homogenized. The final concentration of iodixanol in the photoresin formulation resulted to 15 % w/v. The refractive index of *OptiPrep* was assumed around 1.429.^61^ Mixing was ensured by vortexing the solution after all components were added. The formulation was left at 45 °C until mixing-induced bubbles were removed and filled in single-use, UV-transparent cuvettes *(Carl Roth GmbH, Germany)*. The cuvettes were kept light-shielded and left at elevated temperatures to remove potential bubbles from pipetting. Once no remaining bubbles were observed, cuvettes were moved to a fridge and the photoresin was left to gelate for 15 min. Samples were printed immediately after. The refractive index of the cell-free photoresin was determined with the Abbe refractometer *2WAJ (OPTIKA S.r.l., Ponteranica, Italy)*, calibrated with UPW. Where indicated, 4-hydroxy-TEMPO *(Sigma-Aldrich, USA)* was added at a concentration of 5 ppm. For the Biobooster formulation, the base formulation was prepared as described above, with the addition of TEMPO-OH and Biobooster ( provided by *xolo GmbH, Germany)*. A 50 mM stock solution of Biobooster was prepared in UPW with one molar equivalent of NaOH (50 mM) and stored short-term at 4 °C, protected from light. The dissolved Biobooster was added to the GelMA solution at a final concentration of 5 mM. The final Biobooster formulation contained 10 % w/v GelMA, 15 % w/v iodixanol, 5 ppm TEMPO-OH, 100 ppm DCPI, and 5 mM Biobooster.

### 4.2. Grayscaled Xolography

The trabecular bone structure used was taken from *Thingiverse* based on work of Gross et al. (https://www.thingiverse.com/thing:3609648) and transformed into single slices (resolution 3,840 × 2,160 px, print dimensions of 17.778 × 10.000 × 9.000 mm^3^) with a layer height of 5 µm with *Chitubox (Version 1.9.5, Shenzhen CBD Technology Co., Ltd., China)*.^62^ Grayscaled projections were obtained by creating a rectangular bounding box around the trabecular structure with PowerPoint Version 2406 and a custom *Python* script *(Python version 3.13,*

*Python Software Foundation, USA)*

Utilizing the *VideoCreator* script *(xolo GmbH, Germany)*, videos were created from single slices and uploaded to the *xolid* software *(xolo GmbH, Germany)* on the printer. All the other simplified grayscaled geometries were designed using *PowerPoint (Version 2406, Microsoft Corporation, USA)*.

All samples printed with the base formulation were printed using a *Xube*^2^ *(xolo GmbH, Germany)* printer at a speed (*v*) of 3 mm min*^−^*^1^ and violet (405 nm) light-sheet intensity (*I*) of 9 mW mm*^−^*^2^, resulting in an energy dose (*E*) of 9 mJ mm*^−^*^2^, calculated with *E* = *I* · *d/v* and a nominal light-sheet thickness of *d* = 50 *µ*m. For the Biobooster formulation, a speed of 3 mm min*^−^*^1^ and a light-sheet intensity of 10 mW mm*^−^*^2^ were used. For printing, the material’s refractive index parameter was specified as 1.38, and samples were printed at a position of 5.5 mm to reduce the penetration depth and thus scattering of the visible light projection. The violet light sheet was activated 0.1 s prior to printing and remained activated for 0.1 s after printing. To save material, two geometries were printed in each cuvette, providing at least 5 min between prints for full DCPI deactivation and finished polymerization. For the second print in the same cuvette, cuvettes were rotated by 180°, and the projected images of non-symmetrical geometries were mirrored to minimize the potential impact of crosslinking densities in previously printed grayscaled regions. Following printing, samples were kept in the dark for 5 min and then warmed to 37 °C and poured into well plates containing pre-warmed PBS (cell-free samples) or cell culture media (cell-laden samples) and washed until no visible residues of uncrosslinked photoresin were observed. Samples were incubated in fresh PBS or media at 37 °C for an hour, after which PBS or media was replaced. The next day, samples were immersed in 0.1 % w/v lithium phenyl-2,4,6-trimethylbenzoylphosphinate (LAP) in PBS for 15 min, followed by 3 min UV exposure with a curing lamp *(SUN X11 max, Amazon, USA)*. After post-curing, samples were washed once and immersed in PBS for storage or media for ongoing cell culture.

### 4.3. Optimization of minimum free-floating feature size

To establish viable printing parameters for localization-driven fabrication with the base formulation, printable feature size was characterized as a function of scan speed and violet-light dose by Schlieren imaging. Five settings were evaluated per cuvette, with 5 min of DCPI deactivation between exposures. Starting from the previously established baseline (speed 3 mm min*^−^*^1^, intensity 9 mW mm*^−^*^2^), the scan speed was reduced stepwise to determine the resolution limit (Figure S14 E). After base formulation with 4-hydroxy-TEMPO was introduced, the print window was re-established at a fixed scan speed of 1.5 mm min*^−^*^1^ by varying the light sheet energy, again characterized by Schlieren imaging (Figure S14 F).

### 4.4. Cytotoxcity of 4-hydroxy-TEMPO

Cytotoxicity of 4-hydroxy-TEMPO in primary human dermal fibroblasts (HDFs) was assessed following the same procedure reported by Brauer et al., here applied to 4-hydroxy-TEMPO, using HDFs from the same source.^22^ Briefly, HDFs were obtained from human skin biopsies and expanded through outgrowth culture, with tissue collected under grant (EA2/152/24) submitted to the Ethics Committee of the Charité – University Hospital Berlin. All donors provided written informed consent before tissue collection. Viability was evaluated by live/dead staining with calcein-AM and ethidium homodimer-1 and imaged on a *Leica Stellaris 8 (Leica Microsystems GmbH, Germany)* confocal microscope; metabolic activity was assessed with *alamarBlue (Thermo Fisher Scientific, USA)* on a *Tecan Infinite 200Pro plate reader (Tecan Trading AG, Switzerland)*. Cells were exposed to 4-hydroxy-TEMPO at 5 to 1000 µg mL*^−^*^1^ (ppm); live/dead was assessed after 2 h, and metabolic activity after 24 h.

### 4.5. Culture of hMSCs

hMSCs were isolated from human bone marrow of a healthy adult male donor *(1M0125, Lonza, USA; collected under their institutional guidelines and with informed consent; experiments were performed at Eindhoven University of Technology according to Dutch occupational health legislation)* and characterized as previously described.^63^ hMSCs were cryopreserved in fetal bovine serum *(FBS, Serana Europe GmbH, Germany)* containing 10 % dimethyl sulfoxide (DMSO) until use in experiments. Thawed hMSCs (passage 4–6) were collected in high glucose DMEM with l-glutamine *(hg-DMEM, 41966, Gibco, USA)* before seeding (2.5 × 10^3^ cells cm^-2^) and expansion until confluent (7–10 d) in hg-DMEM supplemented with 10 % FBS, 100 U mL^-1^ of Penicillin and 100 U mL^-1^ of streptomycin *(P/S, Gibco, USA)*, 1x MEM Non-Essential Amino Acids *(NEAA, Gibco, USA)* and 1 ng mL^-1^ recombinant human basic fibroblast growth factor *(bFGF, PeproTech, USA)*. Before reaching confluency, hMSCs were dissociated using 0.05 % Trypsin-EDTA *(Gibco, USA)* before inactivating the trypsin with medium containing 10 % FBS. Cells were pelleted (270 x G, 7 min), counted, and resuspended in DMEM *FluoroBrite (Gibco, USA)* and added to the photoresin to yield a final concentration of 1 × 10^6^ cells ml^-1^.

Small MSC spheroids were formed from Y201 MSCs.^57^ Y201 MSCs were thawed as previously described and expanded in 90 % v/v DMEM, 10 % v/v FBS and 1 % Penicillin streptomycin. Spheroids were formed two days prior to printing. Y201 MSCs were dissociated with 0.05 % Trypsin-EDTA and added with a density of 100 cells per microwell to the *AggreWell 400 (STEMCELL Technologies, Canada)* which was previously rinsed with *Anti-Adherence Rinsing Solution (STEMCELL Technologies, Canada)*. For harvesting, aggregates were washed and dislodged with PBS and collected in a 15 ml tube, both pipette tips and tubes were coated with 2 % FBS in PBS to prevent them from sticking to plastic walls. PBS was exchanged by DMEM *FluoroBrite (Gibco, USA)* and added to the prepared photoresin formulation.

Large MSC spheroids were formed from the hMSCs described earlier. Spheroids were formed nine days prior to printing. hBM-MSCs were added with a density of 250 cells per microwell to the 96-well *Gri3D hydrogel microcavity plate (insphero, Switzerland)*. hBM-MSCs spheroids were cultured in a 1:1 (v/v) mixture of aggregate medium as previously described and endothelial medium (prepared from the *EGM-2 BulletKit (CC-3162, Lonza, USA)*) and consisted of EBM-2 supplemented with the kit-provided *SingleQuots* (hEGF, VEGF, R3-IGF-1, ascorbic acid, hydrocortisone, hFGF-*β*, heparin and gentamicin/amphotericin-B) at the manufacturer-specified volumes.^20^ The fetal bovine serum component was omitted. Spheroids were dislodged with PBS, collected using 5 % BSA-coated tips and tubes, pelleted (300 g, 3 min), and added to the photoresin formulation.

### 4.6. Isolation and culture of hPACs

Human primary articular chondrocytes (hPACs) were isolated from anonymized redundant articular cartilage collected during total knee replacement surgery at Máxima Medical Center. The use of redundant tissue was not subjected to the Medical Research Involving Human Subjects Act and was approved by the Local Research Committee *(Medical Review Ethics Committee Máxima MC, no. N16.148)*, as previously reported.^64^ The hPACs were cryopreserved in FBS containing 10 % DMSO until use in experiments. The hPACs were isolated from the cartilage by enzymatic digestion adapted from Kleuskens et al. with 0.2 % collagenase 2 *(Worthington Biochemical Corporation)* and no hyaluronidase.^65^ Thawed hPACs were passaged until passage 3 with hPAC expansion medium (high glucose Dulbecco’s modified Eagle medium *(hgDMEM, 31966; Gibco, USA)*, 10% FBS, 1 % penicillin/streptomycin *(P/S; Lonza, Switzerland)*, 10 mM 4-(2-hydroxyethyl)-1-piperazineethanesulfonic acid *(HEPES, 15630080; Gibco, USA)*, 1 ng/ml transforming growth factor *β*1 *(TGF* − *β*1*, 100-21; Peprotech, USA)*, 5 ng/ml bovine fibroblast growth factor *(bFGF, 233-FB; Peprotech, UK)*) at 37 °C, 5% *CO*_2_ and 5% *O*_2_. Spheroids were formed two weeks prior to printing. Cells were pelleted (350 x G, 5 min), then hPACs were added at a density of either 25,000 or 50,000 cells per well to 96-well U-bottom plates (Greiner Bio-One B.V., The Netherlands), which were previously rinsed with *anti-adherence rinsing solution* (STEMCELL Technologies, Canada). Cells were resuspended in chondrogenic medium (hgDMEM, 5% FBS *(FBS, Gibco)*, 1% P/S *(Lonza, Switzerland)*, 1% L-proline *(P5607, Sigma-Aldrich)*, 1% HEPES, 1% NEAA, 1% *Insulin/transferrin/selenium-plus (ITS+ premix, 354352; Corning, The Netherlands)*, 1% L-ascorbic acid 2-phosphate *(AA, A8960; Sigma-Aldrich, The Netherlands)*, 10 ng ml*^−^*1 transforming growth factor *β*1 *(TGF* − *β*1*, 100-21; Peprotech)*, 0.1 *µM* dexamethasone *(D4902; Sigma-Aldrich)*). Spheroids were washed with PBS and collected using wide-bore pipette tips, then transferred directly into the photoresin formulation at a seeding density of 1.9 spheroids mL*^−^*^1^ (3 per 1.5 mL cuvette).

### 4.7. Culture of printed scaffolds

Printed scaffolds (diameter = 4–6 mm, height = 3 mm) were cultured in four different conditions: i) a full media composition including supplementation for both adipogenic and osteogenic differentiation (full, F), the negative control of a ii) cell culture medium without further supplementation (control, C), and the positive controls of iii) media including osteogenic differentiation factors (osteogenic, O) and iv) media containing adipogenic differentiation factors (adipogenic, A). Differentiation media were based on protocols described by Huebsch et al.^11^ Therefore, adipogenic supplementation was cycled between media changes with two different dexamethasone concentrations, resulting in two reduced media supplementations, F*_red_* and A*_red_*. Supplements were added from stock solutions based on UPW (ascorbic acid, *β*-glycerol phosphate), methanol (dexamethasone), ethanol (3-isobutyl-1-methylxanthine), and DMSO (indomethacine).

### 4.8. Brillouin measurements

A VIPA-based *Brillouin Hyperfine Spectrometer (HF-8999, LightMachinery, Inc., Canada)* with a spectral resolution of 0.6 GHz in combination with a a 660 nm laser *(Cobolt 05-01, Hübner Photonics, Germany)*, a *Nikon Eclipse Ti2-E microscope (Nikon Corporation, Japan)*, and an *ORCA Fusion C14440-20UP (Hamamatsu Photonics, Japan)* high-resolution camera were used for measurements. Pump killer suppression (≈ 60 dB) was optimized with the

*SpektraLok* software *LightMachinery, Inc., Canada)*. Spectrometer optimization and image acquisition were performed in the LabView-based software *SW-11330 (Version 6989, LightMachinery, Inc., Canada)*. Hydrogels were placed in 35 mm glass Petri dishes for repeated measurements of cell-laden hydrogels under sterile conditions. To maintain humidity during imaging, droplets of PBS (cell-free samples) or cell culture media without growth factor supplementation (cell-laden samples) were added on top of the hydrogels, and the Petri dish was sealed with Parafilm. Brillouin maps were acquired by sampling the specimens at room temperature with a 20×/0.45 Plan Fluor objective (*Nikon Corporation, Japan*, resulting laser spot size *<* 3 µm), an exposure time of 200 ms, and a laser power of 40 mW at the sample location. To reduce back reflections, measurements were performed at a distance of 500 µm from the glass. Brillouin shifts and linewidths were reported as extracted by the LightMachinery software. The mean of different regions within spheroid-laden prints was calculated with an analysis script available in the form of a Jupyter notebook.^66^ Pixels in the transition zone between stiff and soft have been excluded from analysis when comparing full intensity with reduced intensity.

### 4.9. 3D registration and voxel-wise comparison

The Brillouin stack was scaled to its known voxel size and registered to the projection input in Python 3.13 with SimpleITK 2.5. A search over depth, in-plane position, and rotation was followed by gradient-descent refinement, maximizing mutual information. Each Brillouin slice was split into stiff and soft regions with Otsu’s threshold (scikit-image 0.26). The registered input was blurred to Brillouin resolution and thresholded the same way. Each voxel was then classified as matched stiff, matched soft, false stiff or false soft.

### 4.10. Nanoindentation

2D stiffness measurements were conducted with a *Chiaro* nanoindenter *(Optics11 Life BV, Netherlands)*. Printed samples were fixed with 4 % w/v agarose (*UltraPure Agarose, Invitrogen, USA*). To this end, agarose was added to pure water, and the mixture was heated until dissolved. A thin layer of agarose was poured into a plastic Petri dish, and the printed, gently blotted dry sample was added on top shortly before the agarose started to solidify. The sample was then solidified in the fridge and rehydrated with PBS once the agarose turned opaque. Samples were stored in the fridge but left at room temperature for at least an hour prior to testing, to reach a temperature equilibrium. For nanoindentation, a spherical probe with a tip radius of 28 µm and a spring constant of 0.44 N m^-1^ or 0.53 N m^-1^ was used. Measurements were conducted in PBS. To capture both grayscaled areas in the gel, we first observed the initial position by brightfield microscopy in the area with lower exposed intensity, then indented along the direction toward areas of full exposure. A grid of 3 × 30 displacement-controlled indentations was acquired (z-above surface of 15 µm, indentation depth of 15 µm and indentation speed of 7.5 µm s*^−^*^1^) with a distance of 100 µm between indentations, covering an area of 200 × 2900 µm. All collected curves were preprocessed and analyzed with a custom-made graphical user interface.^67–69^ In brief, pre-processing was conducted with *NanoPrepare* software to exclude curves with no stable baseline. Remaining curves were processed with the *SoftMech* graphical user interface, where the *Threshold contact point* algorithm was applied, and a Hertzian contact model was fitted to the loading curve up to an indentation depth of 10 % of the probe radius (2.8 µm). From this, the Effective Young’s modulus was calculated.

### 4.11. Sol fraction, macromer fraction and mass swelling ratio

Cylinders (diameter 2 mm, height 4 mm) were printed at grayscale levels of 100, 150, 200 and 255 without post-curing. To assess the effect of post-processing, cylinders printed at grayscale 100 and 255 were either not post-cured, post-cured for 3 min in 0.1 % (w/v) LAP, or washed overnight in PBS prior to the same post-curing step. Each sample was placed in a pre-weighed tube and weighed directly after printing (*m*_wet_*_,t_*_₀_), freeze-dried *(FreeZone 2.5 L, Labconco, USA)* for 24 h and weighed (*m*_dry,*t*0_), incubated in PBS at 37 °C for 24 h and weighed (*m*_wet_*_,t_*_₁_), and freeze-dried again (*m*_dry_*_,t_*_₁_). The sol fraction, macromer fraction, and mass swelling ratio *q* were calculated as:

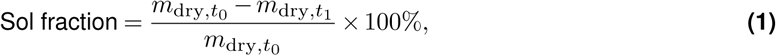

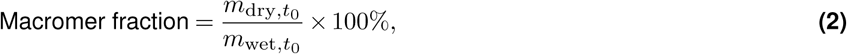

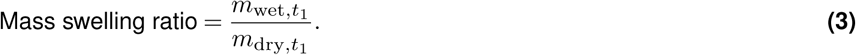

Samples with physically impossible values (negative sol fraction or mass) were excluded.

### 4.12. Fluorescent imaging and image analysis

To assess morphology in grayscaled and fully exposed regions, printed cell-laden hydrogels were sacrificed at different time points and fluorescently imaged. Cellular actin was stained with Alexa 488-conjugated or Alexa 647– conjugated Phalloidin and nuclei with DAPI. At the given time point, samples were washed twice with pre-warmed PBS and fixed with 4 % PFA for 1 hour. PFA incubation as well as all following staining incubations were performed light-shielded on a rocking plate at room temperature. After two washes with PBS, samples were incubated with 1.5 mg ml*^−^*^1^ glycine in UPW for 20 min, permeabilized with 0.3 % Triton for 20 min, and blocked with BSA for 1 hour.

After three washes of 3 min each with 0.1 % Triton, phalloidin (1:250) and DAPI (1:500) were added in 1 % BSA and incubated for 3 hours. Samples were washed three times with PBS and stored in PBS in the fridge until image acquisition. To assess differentiation behavior of hMSCs, calcium was stained with tetracycline hydrochloride and lipid droplets with HCS LipidTox Red, counterstained with DAPI, following a protocol from Vaghi et al. Tetracycline was dissolved in PBS and added at a concentration of 20 µg ml*^−^*^1^ to the cell culture media one day before fixing.^53^

After overnight incubation, samples were washed twice with pre-warmed PBS and subsequently fixed as described above. Samples were then incubated with glycine for 20 min. Following the suggestions of the LipidTox supplier, no permeabilization with Triton was conducted. Instead, LipidTox (1:5000) and DAPI (1:500) were added in 1 % BSA and incubated for 2.5–3 hours. Samples were washed three times with PBS and stored in PBS in the fridge until image acquisition.

Scaffolds were moved to glass-bottom Petri dishes and imaged in air with either the confocal *Zeiss LSM800 paired with the inverted Axio Observer Z1/7 system* and the *ZEN 2.6* (blue edition) software, using a *Plan Apochromat 10×/0.3 NA objective (Zeiss, Germany)* (base samples) or with a *Leica TCS SP8X (Leica Microsystems, Wetzlar, Germany)* and the *LAS X software* using an *HCX PL Apo CS2 10×/0.4 NA objective* (Biobooster samples), using the following excitation/emission parameters: DAPI (Ex: 335 nm; Em: 465 nm), Alexa Fluor 488 (Ex: 490 nm; Em: 525 nm), and Alexa Fluor 647 (Ex: 650 nm; Em: 671 nm). Drops of PBS were added on top of the samples to maintain their hydration during image acquisition. 1,024 × 1,024 pixel z-stacks of about 100 µm were acquired with a voxel depth of 5-10 µm. Scans classified as surface scans were obtained within the first 100 µm of the sample; scans used to investigate cellular morphology inside the hydrogel were taken within a 100–200 µm distance from the surface.

Viability of localized and encapsulated spheroids was investigated with live/dead staining (*LIVE/DEAD Viability/Cytotoxicity Kit, Invitrogen, USA*). In short, constructs were washed once with PBS and stained with 5 × 10*^−^*^6^ M calcein AM and 10 × 10*^−^*^6^ M ethidium homodimer in PBS at 37 C for 30 min. Following one washing step in PBS, samples were submerged in PBS and imaged with the *Invitrogen EVOS M3000 Imaging System* (*ThermoFisher Scientific, USA*) and the dedicated fluorescent LED cubes GFP (482/524 nm) and RFP (542/593 nm) at 4× magnification.

### 4.13. CellProfiler

For quantitative analysis and image batch processing, a custom pipeline was set up in *CellProfiler 4.2.8.* Z-stacks of about 100 µm each were maximum projected into 2D images.^70^ The DAPI channel was smoothed with a Gaussian filter and a typical artifact diameter of 2.5. Nuclei were detected as primary objects based on a typical diameter of 10–50 µm and an adaptive, three-class Otsu thresholding method (threshold range 0–1, 50-pixel adaptive window, smoothing scale 3 and threshold correction factor 0.9). Cytoplasm was identified as secondary object following the Watershed method and an adaptive, three-class Otsu thresholding method (threshold range 0–1, 20 pixels adaptive window, smoothing scale 1 and threshold correction factor 0.75). Samples imaged at 100 µm after 14 days in culture showed higher background staining and smaller nuclei. The lower segmentation threshold was therefore adjusted to 0.08–2 for nuclei and 0.1 for actin staining for some samples. To further include smaller nuclei, the accepted range for primary object diameters was increased to 5–50 pixels. Quantitative data in terms of shape and area, as well as merged and colored projections of representative scans, were obtained with the corresponding CellProfiler modules.

### 4.14. Widefield fluoresence imaging

Fluorescence imaging was performed on a *Zeiss Axio Observer 7 inverted microscope (Carl Zeiss Microscopy, Germany)* using either an *LD Plan-Neofluar 20×/0.4 NA Korr M27* objective for base samples or an EC Plan-Neofluar 5×/0.16 NA objective for Biobooster samples. Channels were acquired using a quad-band beamsplitter (405/493/575/653 nm) with the following excitation/emission parameters: DAPI (Ex: 335 nm; Em: 465 nm), Alexa Fluor 488 (Ex: 490 nm; Em: 525 nm), and HCS LipidTOX Red (Ex: 582 nm; Em: 616 nm). 72-tile mosaics (9 × 8 grid) for base samples or 9- or 12-tile mosaics (3 x 3 or 3 x 4 grid) for Biobooster samples with 10% overlap were stitched and captured as Z-stacks of about 100 *µm* depths.

### 4.15. Widefield fluorescence analysis pipeline

Widefield images were processed using a custom *Python 3.11* pipeline. Linear blending tile stitching was performed, followed by vignette correction using *BaSiCPy 1.2* for visualization; all quantitative analyses were conducted on raw stitched data, considering the inner part of the scaffold to exclude edge effects on the rim. Z-stacks were compressed into single images via contrast-based extended depth of field projection.^71^ Binary masks corresponding to soft and stiff regions were generated from the grayscale printed geometry, manually aligned, and overlaid onto the acquired fluorescence images. Masks were eroded by 100 µm to exclude the transition zone. Nuclear segmentation was performed on DAPI channels using the pre-trained Nuclei Model in *Cellpose 4.0.8.* Alexa Fluor 488 signal quantification was achieved through total fluorescence intensity measurement.^72^ For HCS LipidTOX Red, background subtraction was implemented using white top-hat morphological filtering (Scikit-image 0.25.2), after which a binary threshold and removal of sub-3-pixel-sized objects allowed lipid quantification normalized to the area of the stiff/soft area.^73^

### 4.16. In-situ imaging module

The imaging system comprised a super-apochromatic 2× long working distance objective *(TL2x-SAP, Thorlabs Inc., USA)* and an achromatic doublet tube lens with f = 75 mm *(AC254-075-AB, Thorlabs Inc., USA)*. Brightfield illumination was provided by the LED-based DMD projector integrated in the Xube^2^ system *(xolo GmbH, Germany)*. Images were acquired using a high-resolution sensor *(Alvium 1800 U-1620C, Allied Vision, Germany)*. This configuration achieved an effective magnification of 0.75×, with a sampling resolution of 3.65 µm, and a depth of field of 43.6 µm. The resulting field of view was 19.5 × 11.1 mm and thereby covered the entire printable volume. To reduce the maximum intensity perceived, 1.0 and 0.6 OD filters *(Thorlabs Inc., USA)* were added between the tube lens and sensor. The imaging system was mounted on a motorized translation stage (8MT173-20, Standa, Lithuania) controlled with a 1-axis stepper controller (8SMC5-USB-B8-1, Standa, Lithuania).

### 4.17. In-situ imaging calibration

Autofocus was performed by translating the imaging system along its full travel distance (20 mm) while the projector displayed a high-spatial-frequency USAF 1951 resolution target, using the variance of the Laplacian of the captured image as the focus metric.^74^ The global focus maximum was located by a golden-section search. For xy calibration, ChArUco boards were used, as their ArUco fiducials provide uniquely identifiable markers while the underlying checkerboard pattern allows high sub-pixel registration accuracy.^75^ The geometric rigid registration mapping (homography) between the camera-captured image and the projected input pattern was estimated by Random Sample Consensus (RANSAC).^76^ Both autofocus and ChArUco calibration were repeated twice per calibration run and averaged to reduce variability. Calibration was rerun before every print session. Because the sensor area exceeded the optically corrected region of the objective, focal distortions occurred. These were corrected computationally by modeling the objective and tube lens in *Zemax OpticStudio 2025 R1 (Ansys Inc., USA)* using the manufacturer-supplied (black-box) lens files (Thorlabs, USA). To validate the simulated field curvature experimentally, a cross-shaped pattern of dots (∅ = 25px, spanning 2000 px across the center) was mapped from image to projector space using the ChArUco calibration and displayed by the projector. The pattern was imaged across the full travel range of the imaging system, and the position at which each dot reached its minimum imaged diameter was taken as the in-focus position. Field curvature was then assessed from the focus position as a function of radial distance from the optical center of the objective.

### 4.18. Localization pipeline

The location-driven workflow starts by translating the cuvette in steps of 100 frames, covering the full 10 mm depth. After each step, the imaging system was repositioned to correct for the focal shift, and a frame was acquired. Particles were detected by combining Canny edge detection with morphological operators (edge dilation and closing) for edge-based boundary detection. Because particles spanned multiple focal planes depending on their size and the z-sampling rate, they were tracked across frames using the Trackpy library, and each track was collapsed to a single 3D coordinate (x,y,z) by computing its mean position.^77^ These coordinates served as input for generating printable geometries: for encapsulation, 3D CAD shapes were imposed onto the detected locations, sliced into individual projection frames with a custom slicer, and exported as videos to *xolid (xolo GmbH, Germany)* for printing. When geometries assigned to adjacent spheroids overlap, the slicer generates the overlapping region by taking the union of both geometries and assigning the highest grayscale value to overlapping pixels.

### 4.19. Localization of different objects

Six experimental groups were prepared to demonstrate object detection across a range of sizes. Polyethylene microspheres (∼*500 µm, Cospheric LLC, USA)* were dispersed at 7 microparticles mL*^−^*^1^. Polystyrene microspheres (106 ± 1.9 *µm, Lot PS-FR-SA42, Micro Particles GmbH, Germany)* and polystyrene microspheres (19.85 ± 0.31 *µm, Lot PS-FR-Fi368, Micro Particles GmbH, Germany)* were dispersed at 19 particles mL*^−^*^1^ and 3,875 particles mL*^−^*^1^, respectively; the latter were imaged only, as print parameter optimization for single cell-scale microparticles was not within the scope of this work. hPAC spheroids (1.9 spheroids mL*^−^*^1^), hMSC spheroids, and Y201-MSC spheroids were cultured as described previously. Imaging and localization took up to 4 min per group, depending on object density, and were directly followed by printing. Accuracy and precision are reported as the sample mean and 1 SD, respectively. Localization was evaluated across a minimum of four independent samples per population, with the exception of the 20 µm synthetic particles and Y201-MSC spheroids, for which one and three samples were assessed, respectively. Encapsulations intersecting with the resin meniscus or too close to the wall were not assessed and were considered outliers.

### 4.20. Detection performance

Large objects (PE, hPAC) were annotated manually in the full image stack, blinded to the pipeline output. For small objects (PS, hMSC), every detection found by only one of the pipeline and an independent reference detector (multi-scale Laplacian-of-Gaussian, no shared code), plus a random 15% of shared detections, was judged by a blinded observer. Detections were matched one-to-one; particles at the walls or with best focus at the stack ends were excluded.

### 4.21. Synthetic microparticles and spheroid encapsulation

All groups were encapsulated in grayscale trapezoid constructs. Polystyrene microspheres (106 ± 1.9 µm) were printed in the 4-hydroxy-TEMPO-containing formulation (1.5 mm min*^−^*^1^, 13 mJ mm*^−^*^2^). All remaining groups were printed in the base photoresin formulation (3 mm min*^−^*^1^, 9 mJ mm*^−^*^2^) without 4-hydroxy-TEMPO. hMSC spheroids and PS 106 µm were segmented with a minimum object diameter of 70 µm to exclude debris; hPAC spheroids and PE 500 µm microspheres were segmented with a minimum diameter of 200 µm. Prior to printing, the resin was cooled and maintained at 20 *^◦^*C for 20 minutes.

### 4.22. Perfusable bioreactor printing

Perfusable constructs were printed using the 4-hydroxy-TEMPO-containing formulation at a print speed of 2 mm min*^−^*^1^ and a UV energy of 10 mJ mm*^−^*^2^. Following printing, constructs were transferred to PBS supplemented with 100 mM NaCl, pre-heated and maintained at 40 C in a water bath, to minimize swelling. Post-curing was performed with LAP as described above. After 30 minutes, the PBS solution was refreshed, and constructs were washed for 2 hours. Constructs were then manually perfused with a UPW-diluted red dye *(Tissue Marking Dye Kit, Sigma Aldrich, USA)*, using a 200 µL pipette.

### 4.23. Schlieren imaging

Schlieren imaging was performed using a two-mirror setup based on the open-source 3D-printed design by Jerrod Hofferth (https://makerworld.com/en/models/987486-schlieren-imaging-kit-visualize-density-changes). Two 114 mm aperture, 900 mm focal length Newtonian reflector mirrors served as the primary optical elements. An *Olight S1R Baton II* flashlight was used as illumination source. A PVC color film was used instead of a razor plate as the Schlieren cutoff element. Images were captured with a *Nikon D700 DSLR* camera equipped with an *AF-S VR-Nikkor* 70–200 mm lens in combination with a *Nikon AF-S TC-20E III 2.0×* teleconverter. All images were acquired at f/5.6, 1/80 s shutter speed, and ISO 500.

### 4.24. Software development

Parts of the analysis software and localization pipeline were developed with the assistance of large language models *(Claude Opus 4.8 and Claude-Sonnet 4.6, Anthropic PBC, USA)*, which were used for debugging and auxiliary code. The authors reviewed all AI-assisted code and take full responsibility for the content of this publication.

### 4.25. Statistical analysis

Statistical analysis was performed with *Prism 10 Version 10.6.1* from GraphPad Software LLC, USA). Results are considered statistically significant with p < 0.05 (*), very significant with p < 0.01 (**), and extremely significant with p < 0.001 (***) or p < 0.0001 (****) throughout the work. Results are considered not significant with p ≥ 0.05 (ns). Data extracted from gradient 1 were not normally distributed and showed significantly different standard deviations. Therefore, to assess differences in gradient N*^o^* 1, a Kruskal–Wallis test with Dunn’s post-hoc analysis was utilized for pairwise comparison. Due to some groups violating normality, the same test was performed for the acellular and cell-laden hydrogels to test resolution. The Brillouin shift was analyzed in two steps (*N* = 3 independent hydrogels per condition). First, the changes over time and differences between stiff and soft regions within each media condition were assessed using a Two-Way ANOVA with repeated measures for each condition, followed by Šidák’s multiple comparisons test. Second, a Three-Way ANOVA with repeated measures was performed, pairwise comparing the different cell-laden conditions to the acellular control, followed by Tukey’s multiple comparisons test. Time and region were treated as repeated measures, as stiff and soft regions were measured within the same hydrogels at every time point. The Geisser–Greenhouse correction was applied to account for possible violations of sphericity. Two-Way ANOVA with repeated measures was performed to assess the calcium deposition, lipid droplet count and cell counts (DAPI) of stiff and soft regions, pairwise comparing the different media conditions for the Biobooster formulation (*N* = 3), followed by Tukey’s multiple comparisons test. The base formulation (*N* = 1) is reported descriptively. The stiff and soft regions were treated as a repeated measure, as they were measured within the same hydrogel. For all ANOVA tests, normality of the residuals was assessed with QQ plots, and equal standard deviations of the samples were assessed with homoscedasticity plots. For lipid droplet counts, the homoscedasticity plot indicated increasing variance with increasing lipid droplet count, and zero values were present; therefore, the lipid data were square-root transformed prior to analysis. For the mechanical comparisons, outliers were identified and removed using the ROUT method (Q = 1%)

### 4.26. Visualization and scientific exploitation

Plots were designed with *Origin 2023b (OriginLab Corporation, USA)*, and schematics with *PowerPoint Version 2509 (Microsoft Corporation, USA)*. 3D renderings were produced using the open-source software *Blender (4.5 LTS, Blender Foundation, NL)* Reproduction of the new logo of Eindhoven University of Technology as well as its schematic pixelation (Figure 2 B) is permitted for this purpose under corresponding copyright conditions and by institutional approval.

## Supporting information

VideoS1_TemporalBFS_ControlMedia

VideoS2_z-Stack_TrabecularBonePrint

VideoS3_Localization_Calibration

VideoS4_Localization_Printing

VideoS5_Perfusion

## Author Contributions

L.E. and T.C.v.H. contributed equally to this work. L.E.: Conceptualization, Methodology, Investigation, Formal Analysis, Writing – original draft, Visualization, Funding acquisition. T.C.v.H.: Conceptualization, Methodology, Investigation, Formal Analysis, Writing – original draft, Visualization. M.A.G.O.: Investigation, Formal Analysis, Supervision, Writing - Reviewing and Editing. T.L.v.A.: Investigation, Formal Analysis, Visualization, Writing - Reviewing and Editing. N.F.K.: Investigation, Writing - Reviewing and Editing. P.G.: Resources (immortalized Y201 MSC cell line). S.H., K.I.: Writing - Reviewing and Editing. L.A.: Methodology, Writing – Reviewing and Editing. M.S-S.: Conceptualization, Methodology, Supervision, Writing – Reviewing and Editing, Funding acquisition. M.C.: Conceptualization, Methodology, Supervision, Writing – Reviewing and Editing, Funding acquisition.

## Supporting Information

Supporting Information is available on Bioarxiv or upon reasonable request from the authors.

## Acknowledgments

The authors thank Marcus Reuter for his support and advice during hardware modifications. The authors thank Erik Brauer for performing 4-hydroxy-TEMPO cytotoxicity assays. The authors thank Rob Janssen and Simon Kahlenberg for providing and preparing hPAC spheroids, Joël Bovens for the preparation of hMSC spheroids, and Mireia Valero Puigdoènech for guiding the preparation of Y201 MSC spheroids.

The authors acknowledge funding from the Institute for Complex Molecular Systems. L.S. gratefully acknowledges support from the Young Talent Incentives Award from Gravitation Program “Materials Driven Regeneration”, funded by the Netherlands Organization for Scientific Research (024.003.013). M.C. acknowledges funding by the European Union (ERC CoG, InfiniBlood, 101231661). Views and opinions expressed are however those of the author(s) only and do not necessarily reflect those of the European Union or the European Research Council. Neither the European Union nor the granting authority can be held responsible for them. M.S-S. acknowledges financial support from the European Research Council from ERC AdG (Devise, 101054728) and ERC PoC grant (Factorink, 101248403). IBEC is a recipient of a Severo Ochoa Award of Excellence from MINCIN and member of CERCA Programme/Generalitat de Catalunya.

The authors used *Writefull for Overleaf (version 2025.68.0, ThinqLab Ltd, United Kingdom)* integration and for language editing and proofreading of the manuscript and *Claude Opus 4.8* and *Claude-Sonnet 4.6 (Anthropic PBC, USA)* for partial language editing.

