## Supplementary Information for "Mechanoscribed Xolography for Multiscale Mechanical Patterning of Cell-laden Hydrogels"

[1]Biomaterial Engineering & Biofabrication, Department of Biomedical Engineering, Eindhoven University of Technology, P.O. Box 513, 5600 MB Eindhoven, The Netherlands; [2]Institute for Complex Molecular Systems (ICMS), Eindhoven University of Technology, P.O. Box 513, 5600 MB Eindhoven, The Netherlands; [3]Institute for Bioengineering of Catalonia (IBEC), The Barcelona Institute for Science and Technology (BIST), Baldiri Reixac 10–12, 08028 Barcelona, Spain; [4]xolo GmbH, Volmerstraße 9B, Berlin, Germany; [5]Department of Biology, University of York, Wentworth Way, York YO10 5DD, United Kingdom; [6]Orthopaedic Biomechanics, Department of Biomedical Engineering, Eindhoven University of Technology, P.O. Box 513, 5600 MB Eindhoven, The Netherlands; [7]Nanoscopy for Nanomedicine, Department of Biomedical Engineering, Eindhoven University of Technology, P.O. Box 513, 5600 MB Eindhoven, The Netherlands; [8]Centre for the Cellular Microenvironment (CeMi), The Advanced Research Centre, University of Glasgow, 11 Chapel Lane, Glasgow G11 6EW, United Kingdom; [9]Institució Catalana de Recerca i Estudis Avançats (ICREA), Passeig Lluís Companys 23, 08010 Barcelona, Spain; *These authors contributed equally to this work.

#### Supplementary Information

##### Supplementary Notes

1 Algorithm complexity and computational speed . . . 7

##### Supplementary Figures

S1 Absorbance of iodixanol (IDX) and DCPI 6001 . . . 2
S2 Projector LED emission as a function of gray value . . . 2
S3 Impact of post-curing on mechanical properties in grayscaled samples . . . 2
S4 Degradation of a cell-laden grayscaled print without post-curing . . . 3
S5 Printable range of gray levels in the material system . . . 3
S6 A grayscaled envelope supports complex hydrogel structures . . . 4
S7 Brillouin frequency shift and linewidth of the gradient print . . . 4
S8 Brillouin shift and nanoindentation of acellular control scaffolds . . . 5
S9 Cell viability in grayscaled constructs . . . 5
S10 Imaging system inside the Xube$^2$ . . . 5
S11 Background correction of in-situ images . . . 6
S12 Detection of 20 µm particles and small MSC spheroids . . . 6
S13 Computation time of segmentation and tracking . . . 7
S14 Effect of 4-hydroxy-TEMPO on print resolution and cytocompatibility . . . 8
S15 Binary and grayscale channel bioreactor designs . . . 9
S16 Viability of encapsulated hPAC spheroids . . . 9

##### Supplementary Tables

S1 Effect of post-processing on sol fraction, macromer fraction and swelling . . . 10
S2 Detection performance of the localization pipeline . . . 10
S3 Pipeline benchmark on standard Xube$^2$ hardware . . . 10

##### Supplementary Videos

S1 Z-stack of BFS maps of the trabecular bone structure . . . 11
S2 Temporal evolution of BFS maps of a control scaffold . . . 11
S3 Workflow of the in-situ alignment procedure . . . 11
S4 Localization-driven printing workflow . . . 11
S5 Perfusion of the non-localized channel bioreactor . . . 11

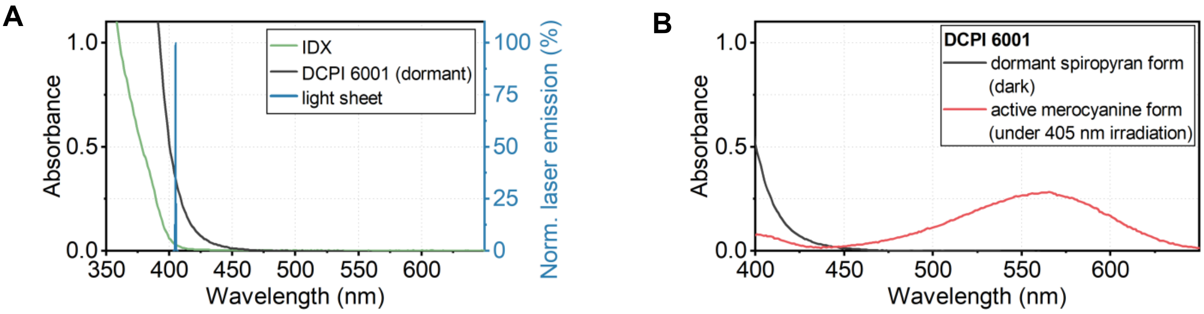


**Figure S1. Absorbance of iodixanol (IDX) and DCPI 6001.** A) IDX shows increasing absorption below 400 nm but negligible absorbance at 405 nm, corresponding to the violet laser emission of the light sheet used in this study. B) The dormant DCPI shows absorption around 400 nm, while the activated DCPI, formed under 405 nm irradiation, shows a characteristic broad absorption band in the range of 450–650 nm. It can therefore be appreciated that DCPI 6001 is the major absorber in this system and penetration depth is not severely affected by the use of IDX.

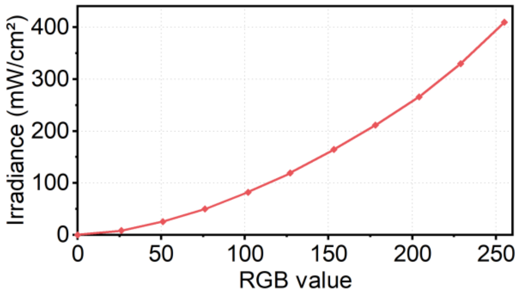


**Figure S2. Emission of Xube² projector LEDs in the focal plane dependent on the applied grayvalue.** Measurement: *Powermeter PM400* and sensor *S130C (both Thorlabs GmbH, Bergkirchen/Lübeck, USA/Germany)* at 565 nm and fixed range 2 W.

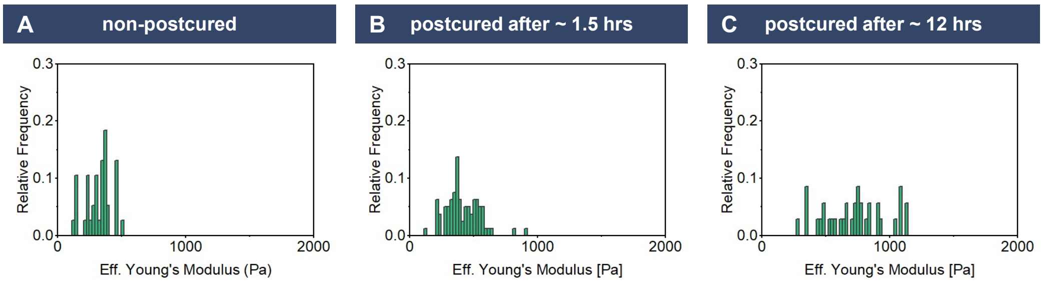


**Figure S3. Impact of post-curing treatment on mechanical properties in a grayscaled sample.** Comparison of the frequency distribution of the effective Young's moduli of a matrix scan covering regions of both full and reduced exposure as measured by nanoindentation. A) Pristine samples show low mechanical properties. B) Low stiffness values occurred less in a grayscaled sample that was post-cured after 1.5 hrs of washing, suggesting that the contrast between grayscaled and non-grayscaled regions diminished. C) Washing of the samples overnight before post-curing increased the overall stiffness slightly and achieved a wide range of stiffness values.

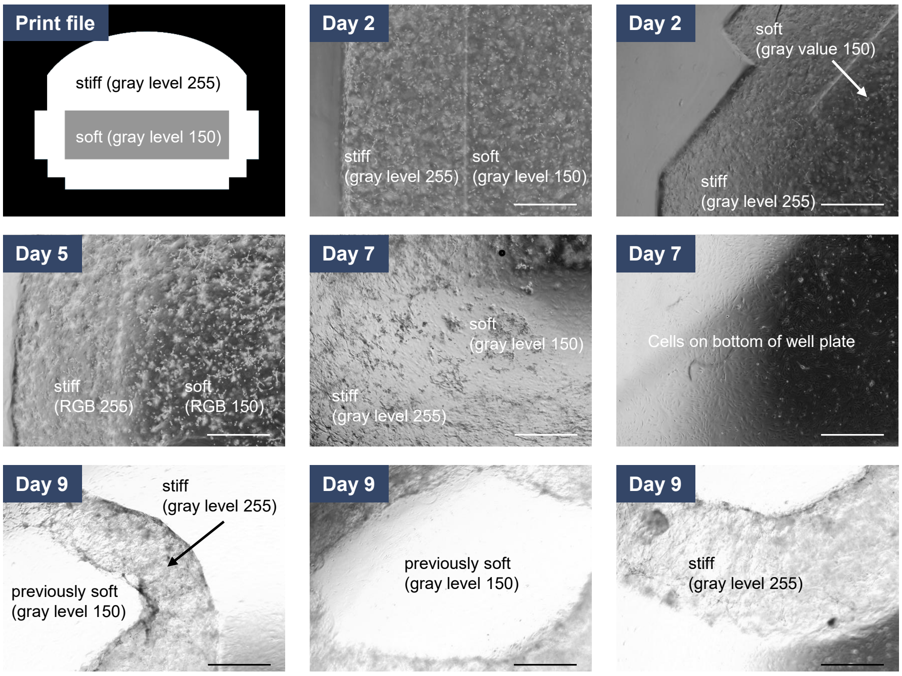


**Figure S4. Test sample to assess hydrogel degradation in a cell-laden and grayscaled print without post-curing.** Prints are based on 10 % w/v GelMA and a cell concentration of $1 \times 10^6$ hMSCs ml$^{-1}$. The print geometry was designed to allow for determining the location of the inner soft region by means of the geometrical outline; however, edges between different gray levels were sufficient to visually distinguish zones. Parts were well defined and showed pronounced edges 2 days after printing. On day 5, degradation was visually observable at the edges of the part. On day 7, a layer of cells was found on the bottom of the tissue culture-treated well plate. The sample area corresponding to a gray level of 150 was entirely degraded after 9 days in culture. Scale bars are 1 mm.

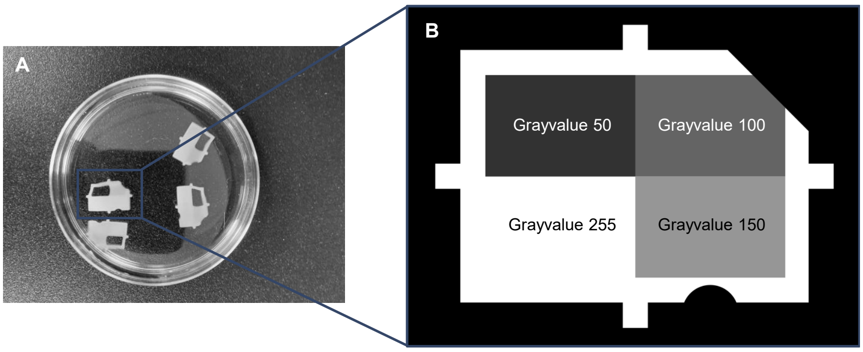


**Figure S5. Test sample to assess the printable range in the utilized material system.** A) Printed part after washing demonstrates that a gray level of 50 is below the threshold to initiate sufficient polymerization. B) Corresponding projection with distinct geometrical features to differentiate different zones.

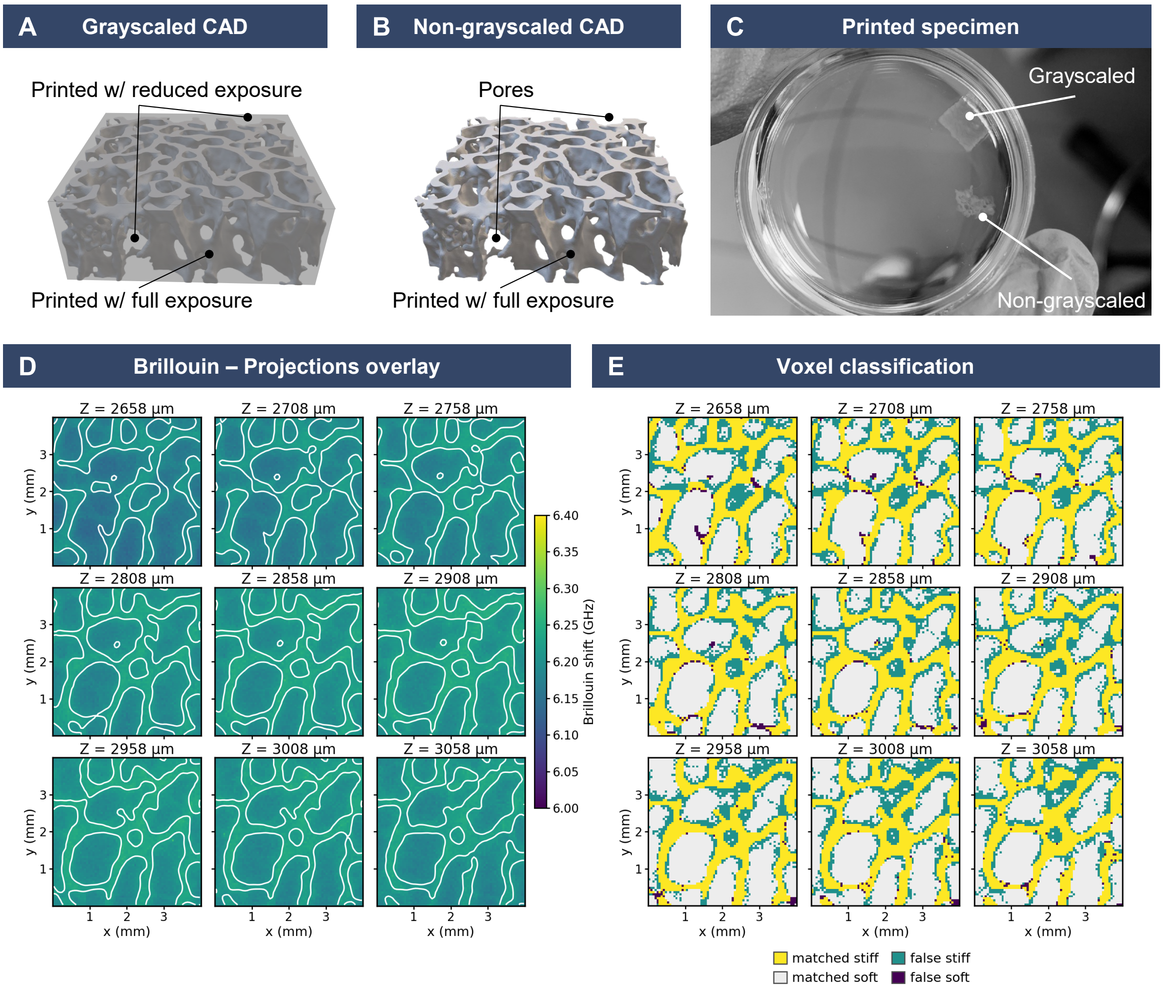


**Figure S6. A grayscaled envelope supports the integrity of complex hydrogel structures.** A) Rendering of the trabecular structure printed at gray level 255, encapsulated in a material envelope crosslinked at a gray level of 100. B) Binary print rendering of the same structure. C) Printed and washed samples demonstrate that the binary-printed samples collapsed due to the thin features and the hydrogel's softness, while the grayscaled part kept its structural integrity. The internal structure of the trabecular remains vaguely visible. D) Brillouin frequency shift maps of all nine slices (50 µm apart), with the stiff/soft boundary of the registered projection overlaid in white. E) Voxel classification of the same slices: Brillouin maps thresholded per slice with Otsu's method and compared with the registered projection (matched stiff, matched soft, false stiff, false soft).

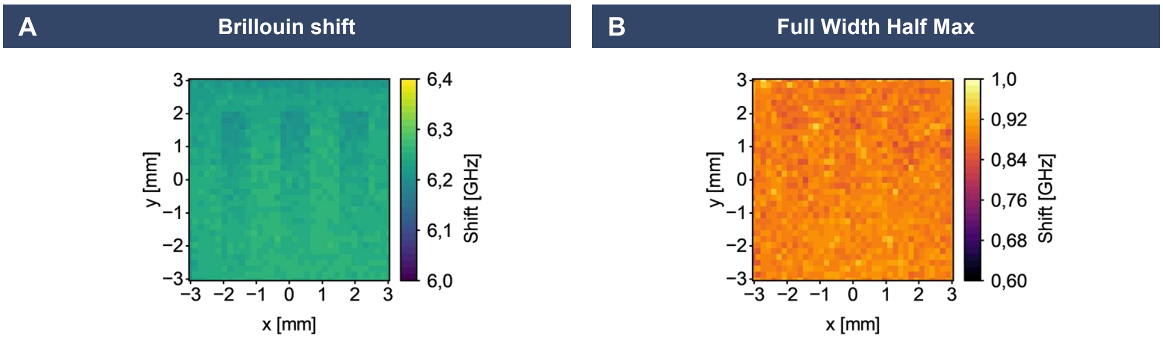


**Figure S7. Brillouin frequency shift and linewidth of the gradient print.** A) Brillouin frequency shift and B) full width at half maximum (FWHM) of a region of the gradient print shown in Figure 2 D. The shift relates to the longitudinal storage modulus, whereas the FWHM (linewidth) reflects acoustic attenuation and is commonly used as a proxy for the viscous (loss) component of the GHz mechanical response. FWHM does not show a large difference between soft and stiff regions.

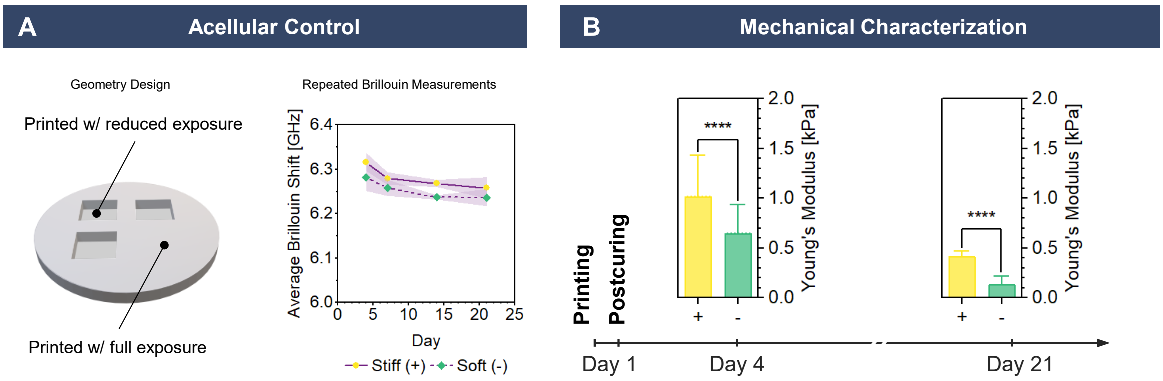


**Figure S8. Acellular control scaffolds.** A) Printed geometry with stiff $(+)$ and grayscaled (gray level 100), soft $(-)$ areas (left) and their Brillouin frequency shift on day 4, 7, 14, and 21 (right). B) Effective Young's modulus of stiff and soft areas on day 4 and 21, measured by nanoindentation. Mean $\pm$ SD, $N = 3$ hydrogels.

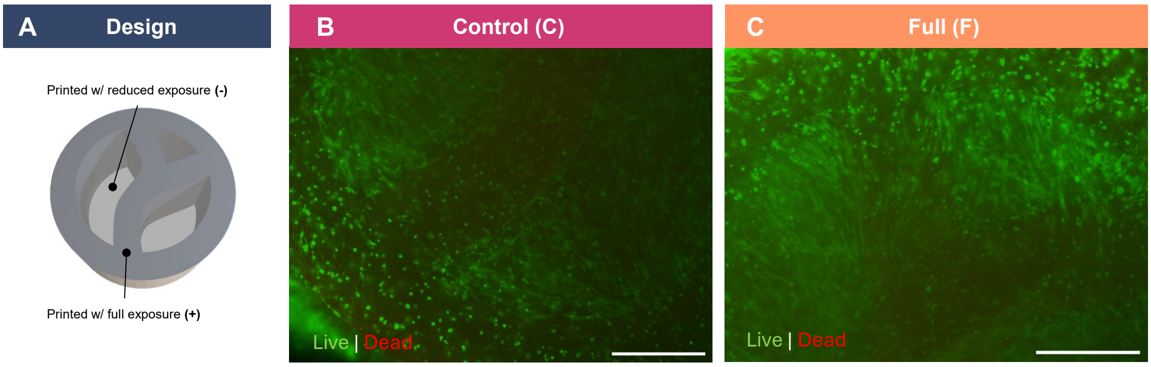


**Figure S9. Cell viability in grayscaled constructs.** A) Design of the simplified trabecular-like structure with stiff and soft regions, as introduced in Figure 2 G. B) Widefield fluorescence images of live/dead staining (live: green, dead: red) of cell-laden samples printed with the BioBooster formulation (Figure 4) on day 21. Scale bars: 1 mm.

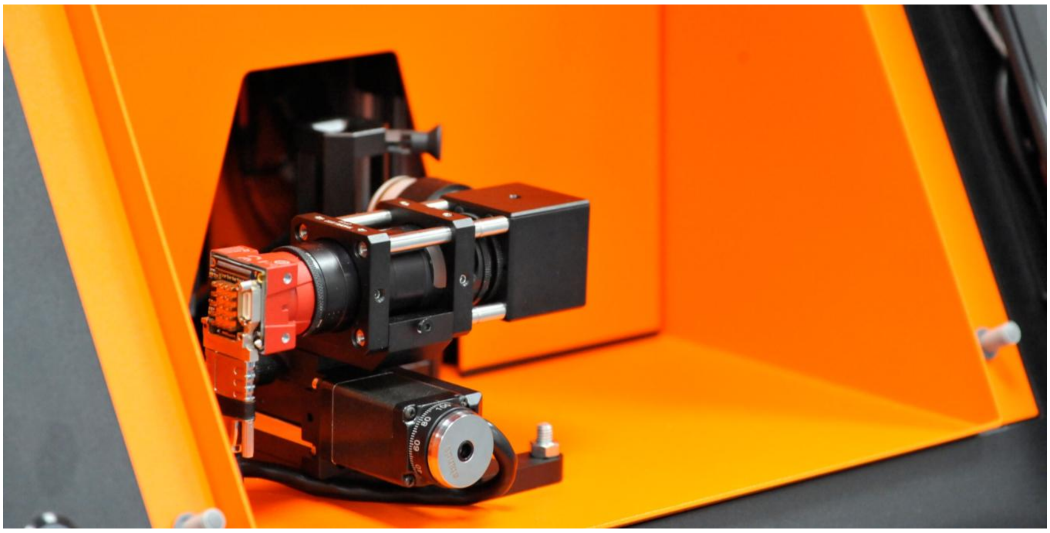

**Figure S10. Photo of imaging system inside Xube$^2$.**

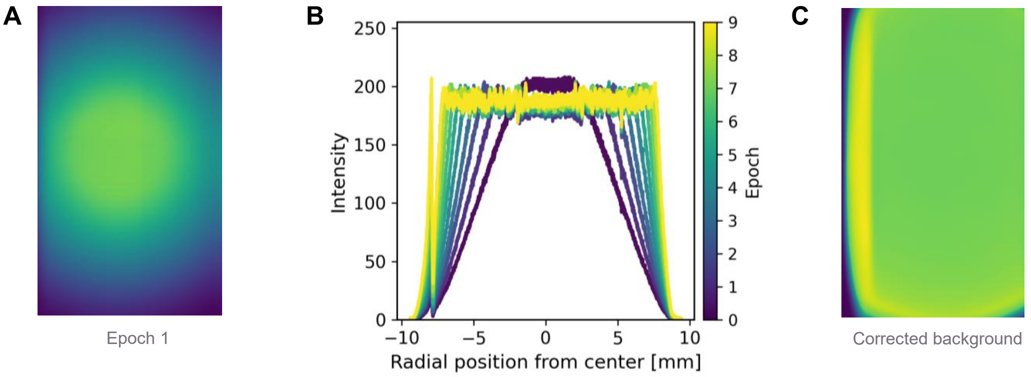


**Figure S11. Background correction of in-situ images.** A) Raw image prior to correction showing non-uniform illumination B) corresponding radial intensity profile C) corrected image with homogeneous background.

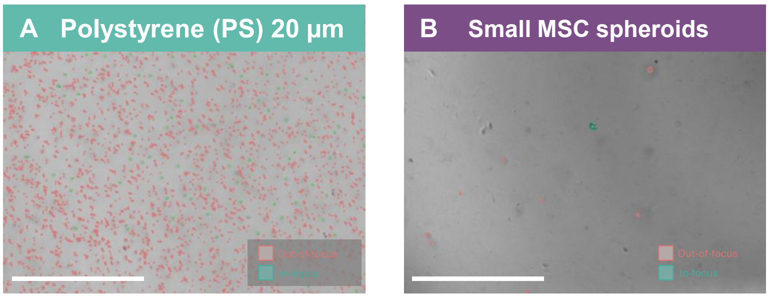


**Figure S12. Detection of 20 µm particles and small MSC spheroids.** Representative images for object detection overlaid on a cuvette, as observed by imaging samples of 20 µm polystyrene particles (A) and small MSC spheroids (B). Scale bars are 2.5 mm.

#### Supplementary Note 1: Algorithm complexity and computational speed

From a theoretical point of view, the particle localization algorithm steps scale differently. Acquisition scales with $\mathcal{O}(N)$, with N the number of frames. Segmentation is linear in N and is dominated by a fixed per-frame cost. The particle-dependent term is linear in the detections per frame. It is the slowest of the image processing steps but is parallelized, so that the per-pixel work scales with the core count. Tracking is $\mathcal{O}(n \log n)$ in the number of detections per frame pair, and quadratic in the candidate links to the previous frame. Resolving particle-to-particle pairs between frames is what produces a computational ceiling. This can be prevented by replacing the frame-to-frame linking with proximity clustering of the full detection cloud, since a z-stack contains no motion that needs to be resolved. Slicing is independent of particle size and linear in the number of localized particles, but cubic in the printed object diameter.

To determine the scalability and computational limitations of the particle localization, a benchmark was built to assess performance on the hardware used in the xolographic printer, a Jetson Xavier NX Developer Kit (NVIDIA, USA). Synthetic stacks provided a computational load at particle numbers beyond the available datasets, so that the pipeline could be timed up to $10^6$ particles/ml. The benchmark generated synthetic data in which spheres of diameter $d_p$, placed at random positions in the imaged volume, were drawn as dark discs on a bright background, blurred in $xy$ by a Gaussian of width $\sigma = \mathrm{NA} \cdot |\Delta z|$, set by the defocus $\Delta z$ of each particle from the imaging plane, faded by $(r/(r+\sigma))^2$ with $r$ the disc radius, and drawn only while $|\Delta z| \leq (d_p + \mathrm{DoF})/2$. The stacks were required only to reproduce the computational load of real data, which was set by the number of frames and the number of detections per frame. When timed against real acquisitions of 104 $\mu$m particles at matched detections per frame, segmentation was at most 28% slower on real data and tracking agreed to within 8%, so the synthetic stacks were a valid basis for timing. The computational time per step for real data runs is reported in Table S3, and Figure S13 shows segmentation and tracking up to $10^6$ particles/ml.

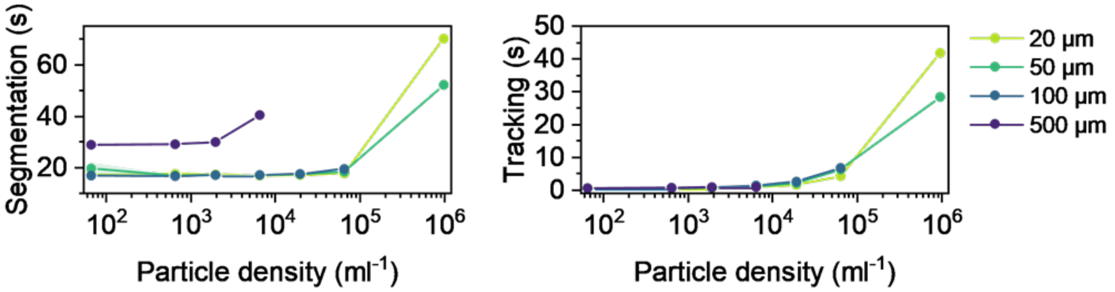


**Figure S13. Computation time of the combined segmentation and tracking steps for synthetic data with different particle densities and particle diameters.** Data are presented as mean ± SD ($N = 3$).

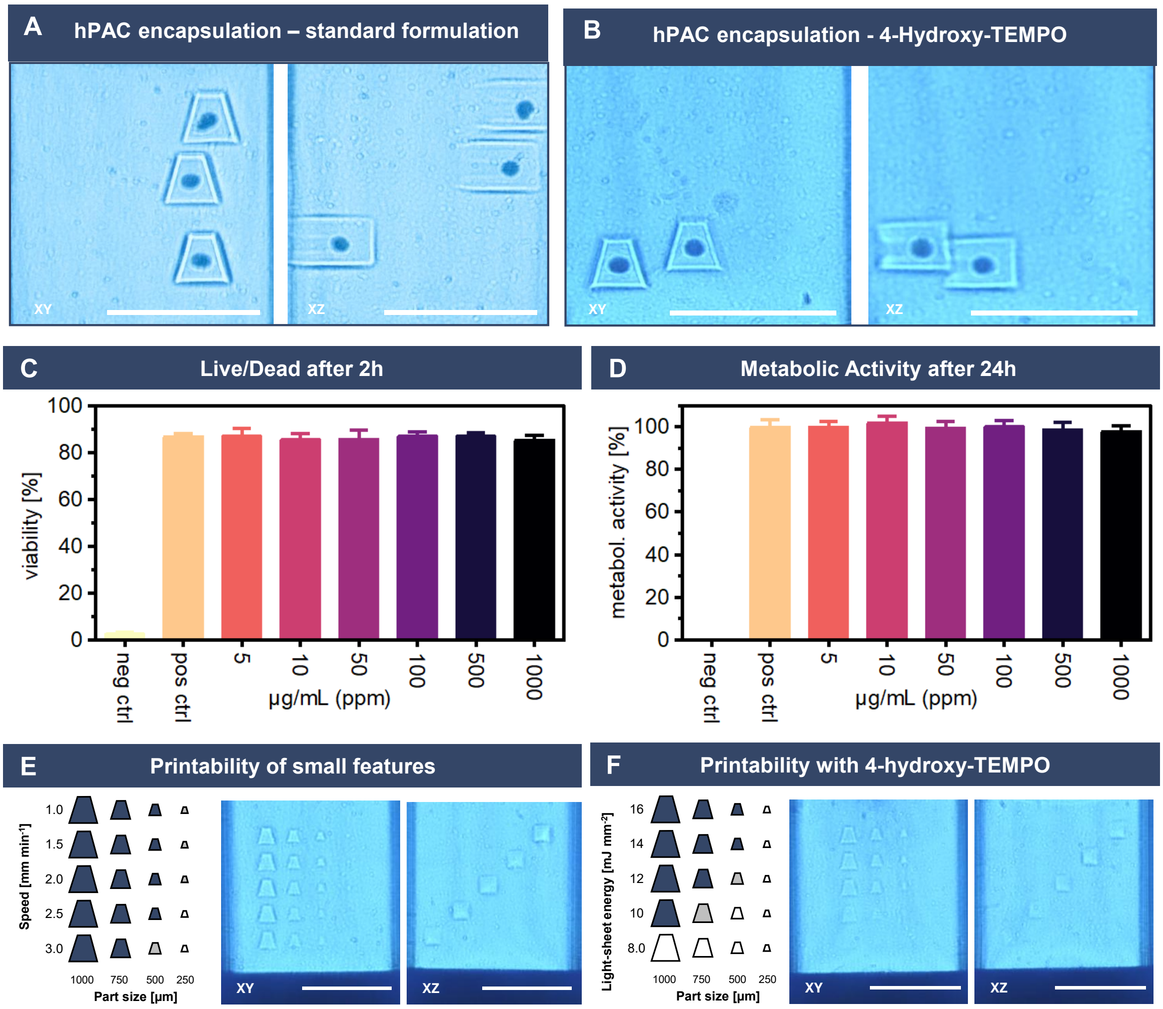


**Figure S14. Effect of 4-hydroxy-TEMPO on print resolution and cytocompatibility.** A) Schlieren image of chondrocyte spheroids printed with the standard formulation. B) Schlieren image of chondrocyte spheroids printed with the TEMPO-containing formulation. Scale bars are 5 mm. C) Viability (% of live cells) of 2D monolayer HDFs after 2 h exposure to 4-hydroxy-TEMPO at 5 to 1000 µg/mL. D) Metabolic activity (normalized to expansion medium control) of 2D monolayer HDFs after 24 h. Data are presented as mean ± SD, n = 12 from two independent experiments. E) Print settings and corresponding Schlieren images of trapezoids within the cuvette by increasing print speed and decreasing print size. F) Print settings and corresponding Schlieren images for material formulation with 4-hydroxy-TEMPO by decreasing energy and decreasing print size.

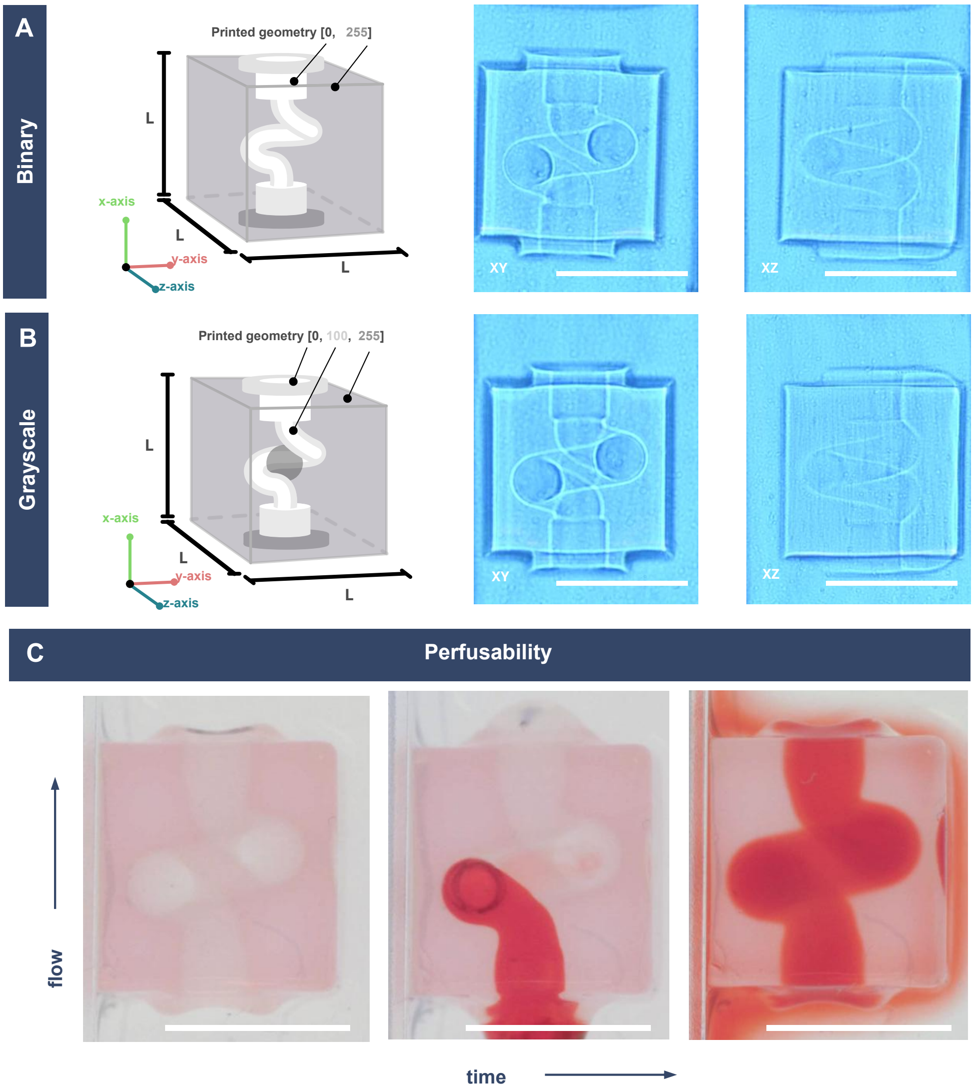


**Figure S15. Non-localized channel bioreactor designs.** CAD models of the binary (top) and grayscale (bottom) non-localized channel bioreactor designs (left, l = 7 mm) with corresponding Schlieren images in the XY and XZ planes. Scale bars: 5 mm.

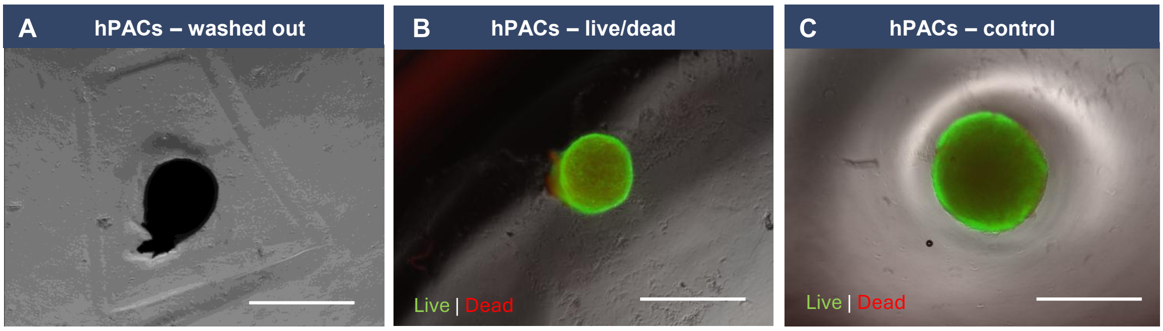


**Figure S16. Viability of encapsulated hPAC spheroids.** A) Brightfield image of encapsulated hPACs spheroids after washing. B) Widefield fluorescence and brightfield overlay of encapsulated hPAC spheroids. C) Widefield fluorescence and brightfield overlay of non-printed hPAC spheroid control. Scale bars: 884 µm.

### Supplementary Tables

**Table S1. Effect of post-processing on sol fraction, macromer fraction and mass swelling ratio of printed GelMA (base formulation).** Postcuring: 3 min in 0.1% (w/v) LAP. Data reported as mean ± SD of $n = 3$ samples per group, except where indicated: [a]$n = 2$, [b]$n = 1$ (see Methods). Low dry sample masses increase the variability, particularly of the sol fraction.

| Grayscale | Post-processing | Sol fraction [%] | Macromer fraction [%] | Mass swelling ratio [–] |
|---|---|---|---|---|
| 100 | None | 68 ± 3[a] | 9.5 ± 1.8 | 11 ± 6 |
| | Postcuring | 37 ± 4 | 8.1 ± 2.4 | 15 ± 3 |
| | Washing + postcuring | 37 ± 6[a] | 3.1 ± 1.1 | 14 ± 2 |
| 255 | None | 67 ± 5 | 9.8 ± 1.0 | 24 ± 5 |
| | Postcuring | 39 ± 17 | 7.6 ± 0.6 | 15 ± 4 |
| | Washing + postcuring | 17[b] | 4.2 ± 0.8 | 12 ± 5 |

**Table S2. Detection performance of the localization pipeline per population.**

| Population | Ground truth[a)] | $N$[b)] | Particles | Detections | FN[c)] | FP[c)] | Recall (%)[d)] | Precision (%)[d)] |
|---|---|---|---|---|---|---|---|---|
| PE, ∼500 µm | Manual | 4 | 31 | 23 | 8 | 0 | 74 (57–86) | 100 (86–100) |
| hPAC spheroids | Manual | 6 | 11 | 8 | 4 | 1 | 64 (35–85) | 88 (53–98) |
| PS, 106 µm | Second detector | 5 | 155 | 155 | 5 | 5 | 97 (93–99) | 97 (93–99) |
| hMSC spheroids | Second detector | 7 | 159 | 159 | 55 | 55 | 65 (58–72) | 65 (58–72) |

[a)]Manual: manual annotation of the full image stack; Second detector: independent second detector with blinded manual checks. [b)]Number of independent samples. [c)]FN, false negatives (missed particles); FP, false positives (detections without a particle). Particles and detections are equal when FN = FP. [d)]95% Wilson score confidence intervals in brackets.

**Table S3. Pipeline benchmark on standard Xube$^2$ hardware.** Each step of the pipeline was tested separately; $N$ denotes the number of independent runs. Benchmarks were performed on synthetic particles of 104 µm diameter, with a density of ∼100 particles per cuvette (1.5 mL). Time is reported as mean ± SD.

| Step | $N$ | Time (s) |
|---|---|---|
| Acquisition | 5 | 122.6 ± 0.5 |
| Segmentation | 5 | 22.2 ± 1.9 |
| Tracking | 5 | 0.69 ± 0.17 |
| Slicing | 3 | 79.1 ± 0.3 |
| Total | | 219 ± 1 |

#### Supplementary Videos

**Video S1. Z-stack of the BFS maps captured from a section of the trabecular bone structure as shown in Figure 2 F.**

**Video S2. Temporal evolution of BFS maps measured on day 4, 7, 14 and 21 for the printed scaffold from the control group shown in Figure 3 which was cultured in maintenance media.**

**Video S3. Workflow of the in-situ alignment procedure.** For z-alignment, a USAF 1951 target was projected while the camera was translated along the optical axis, and the in-focus plane was identified using a golden-section search. For xy-alignment, ChArUco boards were projected at the start and end positions of the cuvette to register the projector to the imaging coordinates.

**Video S4. Printing workflow.** A z-stack was first acquired by translating the cuvette, while the imaging system was also translated to correct for focal shift. A trapezoid encapsulation structure was then printed, and the video visualizes the intersection of the particles with the projected light.

**Video S5. Perfusion of the non-localized channel bioreactor shown in Figure S15 A and C.**
