## Supplementary figures and images for "Mechanoscribed Xolography for Multiscale Mechanical Patterning of Cell-laden Hydrogels"

### VideoS1_TemporalBFS_ControlMedia

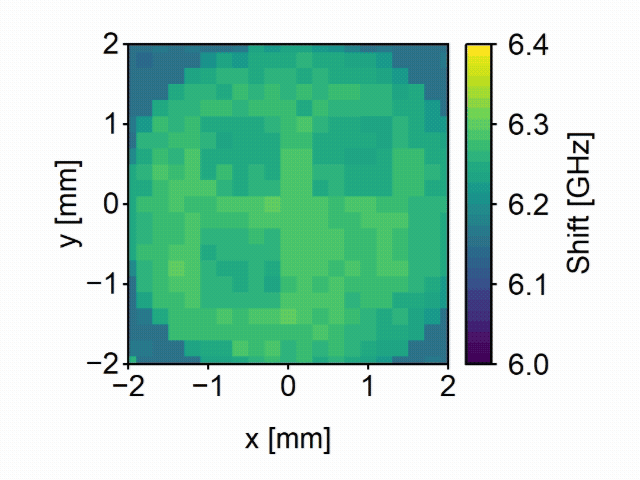

### VideoS2_z-Stack_TrabecularBonePrint

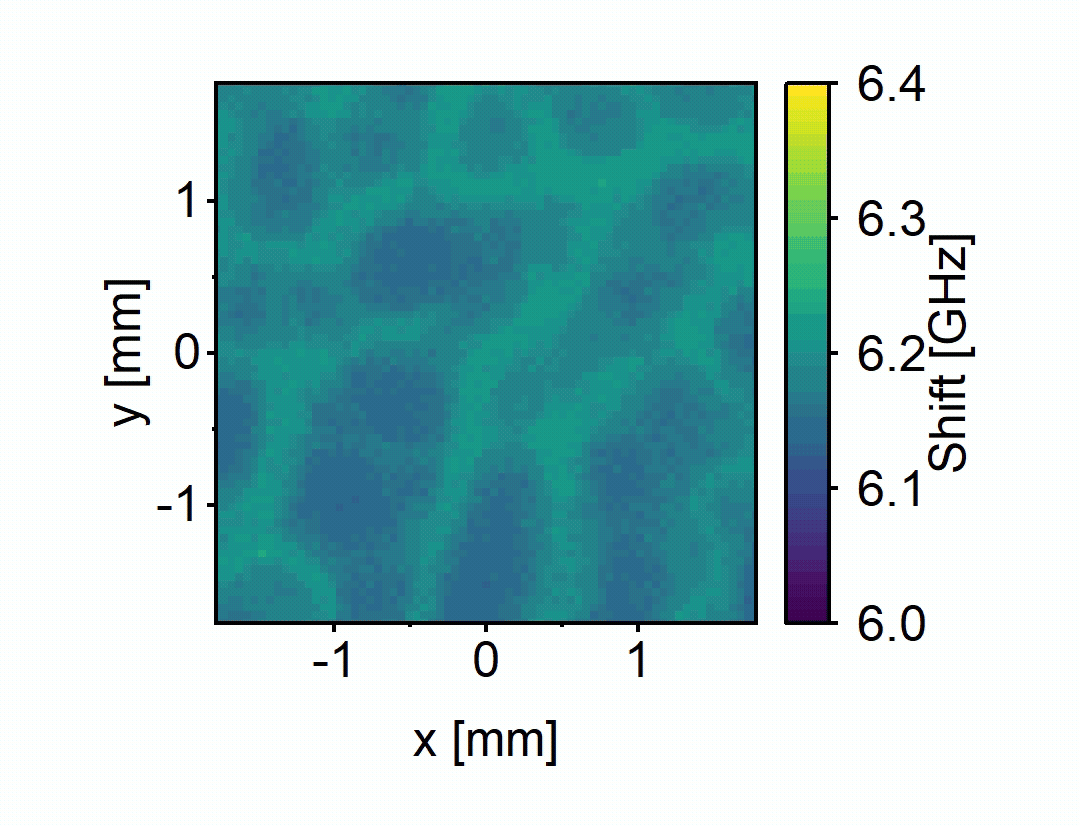
